# Conserved covariance structure underlies 60 million years of morphological diversification in primates

**DOI:** 10.64898/2026.09.09.748908

**Authors:** Anna Penna, Diogo Melo, Bruce S. Martin, Monique N. Simon, Daniela M. Rossoni, Barbara A. Costa, Guilherme Garcia, Felipe B. de Oliveira, Gabriel Marroig, Fabio A. Machado

## Abstract

Evolvability—the capacity of populations to respond to selection—is shaped by the structure and amount of variation transmitted from generation to generation.Whether the heritable covariance structure itself remains stable or evolves rapidly is a central, yet unresolved, question in phenotypic evolution. Competing hypotheses suggest that trait covariation may be constrained by developmental and genetic architectures or, alternatively, reshaped by persistent directional selection. Here, we test whether covariance structure and evolvability are stable over macroevolutionary timescales by applying a comparative evolutionary quantitative genetics framework on primates. We quantified cranial variation using over ten thousand specimens representing 309 species and compared phenotypic covariance matrices for 57 genera within a Bayesian framework. Our results show that despite extensive morphological divergence,primates show remarkably conserved patterns of variation, modular organization, and capacity to respond to selection. Reconstruction of selection gradients across the primate radiation revealed that selection was highly structured, with preferential directions aligned with major axes of cranial variation. These results suggest that the stability of evolvability does not reflect evolutionary stasis, but rather emerges from the alignment between conserved developmental architecture and selection. Our findings demonstrate that covariance structure and the evolutionary potential it provides can persist over deep evolutionary timescales, providing a mechanistic framework for understanding how developmental systems shape the trajectories of major radiations.

---

**D**espite the extraordinary morphological diversification observed across the tree of life, evolutionary change is often biased toward particular directions and combinations of traits. Understanding how such biases emerge and persist over deep evolutionary time is central to the study of evolvability: the capacity of a biological system to generate heritable phenotypic variation that can respond in the direction of selection (Hansen, Houle, et al., 2023; Kirschner and Gerhart, 1998). In classical quantitative genetics terms, evolvability is often operationalized as the amount of additive genetic variation of a trait (Hansen, Pélabon, and Houle, 2011), which ultimately determines the rate of phenotypic evolution and the increase in mean fitness caused by directional selection (Lande, 1976). Over short evolutionary timescales (spanning tens of generations or fewer), evolvability effectively predicts responses to selection and helps identify genetic constraints (Steppan, Phillips, and Houle, 2002; Arnold et al., 2008). However, whether evolvability plays a consistent role in shaping macroevolutionary patterns remains an open question (Uyeda and Machado, 2025; Schluter, 2024; Tsuboi et al., 2024; Machado, Mongle, et al., 2023).

In phenotypes composed of multiple interacting elements (i.e. complex traits), evolvability is closely related to the concept of morphological integration: the degree to which traits co-vary due to developmental, functional, or genetic interactions (Olson and Miller, 1958; Gould, 1985; Snell-Rood and Ehlman, 2023). For example, the bones of the mammalian skull are simultaneously shaped by brain growth, facial development, accommodation of sensory organs, and masticatory function (Hallgrímsson et al., 2009; Moss and Young, 1960; Koyabu, 2023). These various morphological and developmental systems reflect a genetic architecture that leads to differences in the degree of association (integration) and relative independence (modularity) among traits in different regions of the skull (Wagner, 1996). Given that trait association governs the ability of populations to track shifting selective pressures due to correlated response to selection (Lande, 1979), morphological integration is thus an essential component of evolvability (Schluter, 1996; Hansen and Houle, 2008; Marroig, Shirai, et al., 2009; Melo, Porto, et al., 2016).

A key challenge to the notion that evolvability can have macroevolutionary consequences is that the amount and structure of heritable variation, summarized by the additive genetic covariance matrix (G-matrix), can also evolve by drift, mutation, and, more importantly, directional selection (Turelli, 1988; Walsh and Blows, 2009; Pigliucci, 2006; Barnett, Meister, and Rainey, 2025; Jones, Arnold, and Bürger, 2007; Pavlicev, Cheverud, and Wagner, 2011; Melo and Marroig, 2015; Penna et al., 2017; Milocco and Salazar-Ciudad, 2022). This has led to the expectation that genetic (co)variances (and, by extension, evolvability) may change on the timescale of several generations, thereby hampering the use of microevolutionary models to understand macroevolution (Shaw et al., 1995; Jones, Arnold, and Bürger, 2007; Doroszuk et al., 2008). Yet, a number of recent macroevolutionary studies have shown that large-scale phenotypic patterns often align with microevolutionary predictions, suggesting some stability in evolutionary potential across deep time (Houle et al., 2017; McGlothlin et al., 2018; Machado, Mongle, et al., 2023; Rohner and Berger, 2023; Rohner and Berger, 2025; Opedal et al., 2023; Holstad et al., 2024; Simon, Courtois, et al., 2025). Understanding whether these macroevolutionary patterns are underpinned by stable genetic architectures or whether covariation itself evolves is thus crucial for understanding how processes that occur at the population level (microevolution) scale up to influence macroevolutionary trends. Furthermore, because macroevolutionary dynamics are largely governed by directional selection that can change covariance patterns (Simpson, 1944; Lande, 1980; Lande, 1981), it is essential to elucidate the mechanisms by which such stability can arise and be maintained.

To evaluate the long-term stability of evolvability, we examine the structure of cranial trait (co)variance in the third most diverse order of living mammals: Primates, whose exceptional ecological diversity is mirrored by substantial variation in cranial morphology (Fleagle, Gilbert, and Baden, 2010). The evolutionary history of primates is characterized by both rapid radiations and long-term stasis (Herrera, 2017; Arbour and Santana, 2017; Pozzi and Penna, 2022), making them an ideal system for evaluating the stability of trait covariation patterns across a broad phylogenetic scale. Using a Bayesian framework that integrates quantitative genetics and phylogenetic comparative methods, we test whether phenotypic covariance matrices (P-matrices) exhibit long-term stasis or divergence across the primate phylogeny. Although G-matrices are central to theoretical models of evolvability, they remain difficult to estimate in non-model systems. P-matrices provide a practical alternative because, for morphological traits, patterns of phenotypic covariation closely parallel the underlying genetic covariance structure (Cheverud, 1988; Roff, 1997; Steppan, Phillips, and Houle, 2002; Holstad et al., 2024). This correspondence is thought to arise because genetic, mutational, and environmental sources of variation are funneled through shared developmental dynamics to produce similar phenotypic covariance structure, a phenomenon observed across several systems (Houle et al., 2017; Noble, Radersma, and Uller, 2019; Rohner and Berger, 2023; Milocco and Uller, 2026). By quantifying and comparing the structure of P-matrices across 57 primate genera, our study provides an unprecedented large-scale test of whether phenotypic (co)variance and, by extension, the evolutionary potential, remain conserved or evolve over deep time. In doing so, we provide new insights into the persistence of developmental constraints, the evolution of trait integration, and the underlying mechanisms behind the long-term dynamics of evolvability in a major vertebrate radiation.

## Results

### Conserved structure of trait association despite morphological disparity

We examined patterns of morphological variation in cranial shape and size based on 37 linear measurements collected from 10,073 adult specimens representing 309 species Figure 1A). Primates show extraordinary diversity in facial morphology, ranging from dolichocephaly (long skulls with an elongated, narrow rostrum, *e*.*g*. lemurs) to extreme brachycephaly (broad skulls with a flatter, less projecting face, *e*.*g*. humans). Principal component analysis (PCA) revealed a well-structured morphospace dominated by variation along PC1 (86.59%), corresponding to cranial size (*see SI Appendix* Fig. S1A for the regression of PC1 on size). This axis also captures relative orbit size and convergence, as well as the flexion of the cranial base region (Fig. 1C). The second major axis of variation (PC2 5.25%) primarily captures shape, specifically, a contrast between the relative height of the cranial vault and the elongation of the rostrofacial region. The five major clades (Strepsirrhini, Tarsiiformes, Platyrrhini, Cercopithecidae, and Hominoidea) occupied distinct regions of the morphospace, reflecting clade-specific allometries that were nearly parallel to each other along the size gradient. Specimens of *Microcebus, Cebuella, Alouatta*, and *Homo* showed little or no overlap with any other genera, while Cercopithecidea has the highest overlap both within and among clades (Platyrrhini and Hominoidea, in particular with lesser apes). Tarsiiformes are much more similar to Strepsirrhines than to their closer relatives within Haplorrhini. A PCA analysis removing the effects of size also showed minimal overlap between major clades, but less variation concentrated along the main axes of variation (PC1 30.92% and PC2 20.7%; *SI Appendix* Fig. S1C). All further analyses were performed in the original P-matrices, including size.

**Figure 1.**
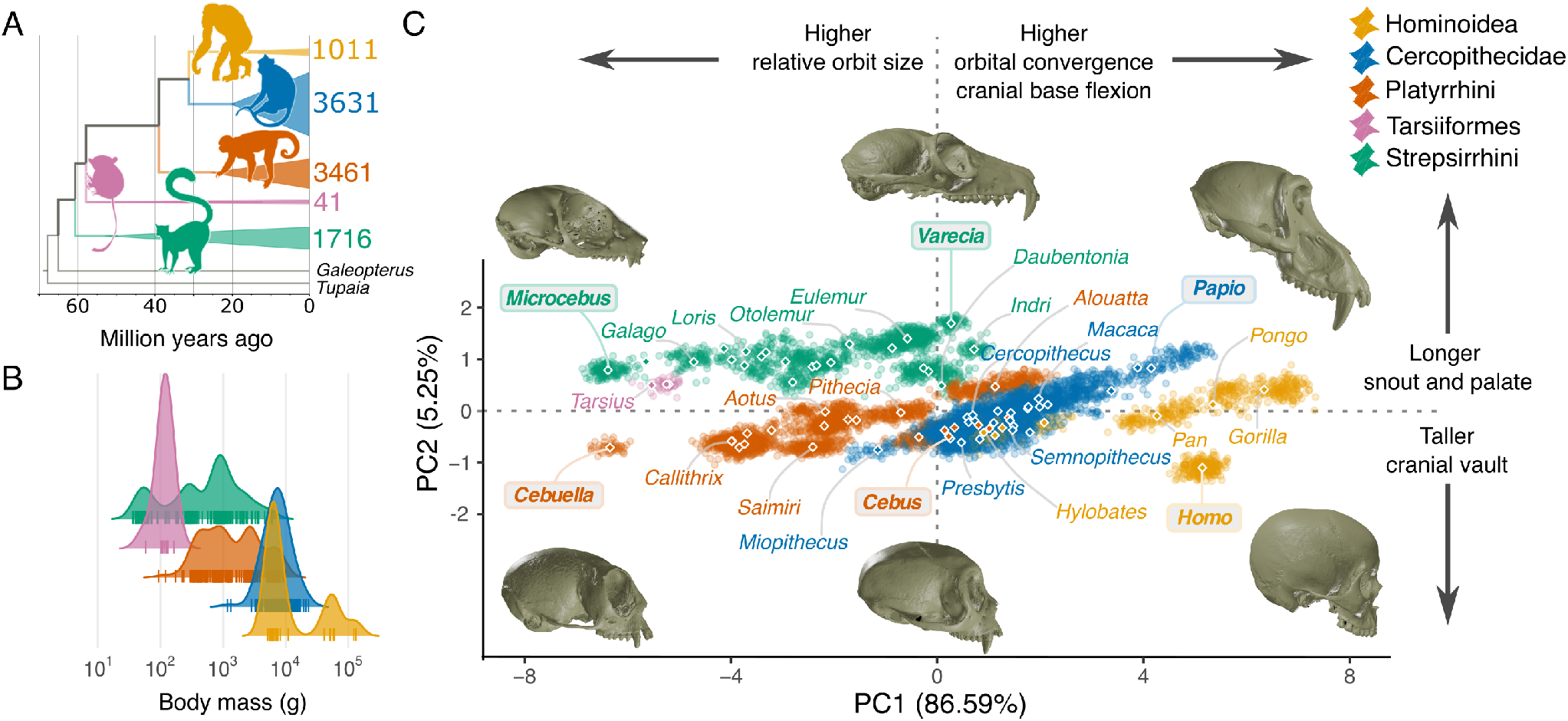
Primate phenotypic diversity across major clades. **(A)** Phylogenetic relationships and divergence times among the five major primate clades (colors), and respective sample sizes in the cranial morphometric dataset. **(B)** Distribution of species average body mass across major primate clades (colors), which varies by five orders of magnitude. Each tick mark represents a living species (N = 419). **(C)** Cranial morphospace based on 37 linear measurements obtained from almost ten thousand specimens, visualized using the first two principal components of a PCA. The percent of variance explained by each axis is shown in parentheses. Points represent individual specimens; diamonds indicate genus-level centroids. Crania for six selected genera (highlighted labels) showcase the broad range of morphological variation in facial length, encephalization, and flexion of the cranial base. Silhouettes downloaded from phylopic.org (CC0 1.0 Universal Public Domain Dedication license and Public Domain Mark 1.0 license); Skulls (not to scale) from MorphoSource, used with permission.

Comparisons of the P-matrices across all primates reveal remarkable stability in trait (co)variance structures (Fig. 2).

**Figure 2.**
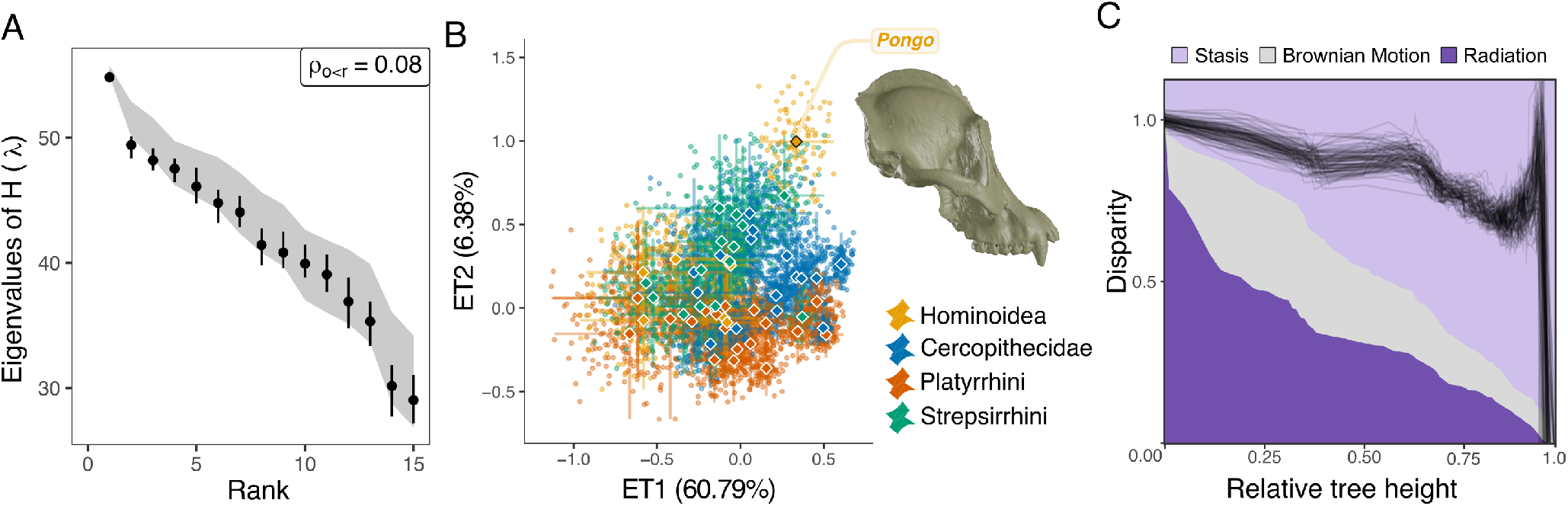
Phenotypic matrices exhibit conserved similarity in geometric structure. (**A**) Comparison of the 57 **P**s using the Krzanowski common subspace. Rank-specific eigenvalues of **H** (λ) from the empirical posterior distribution (black) compared to the null envelope derived from randomized matrices (gray polygon). The dots represent posterior medians, and vertical bars indicate 95% highest posterior density intervals. λ approaching 57 indicate stronger shared subspace alignment across all matrices. Overlap between empirical and null distributions indicates strong alignment in the subspaces, compatible with the null hypothesis that genus-specific covariance structure is not required to explain the observed pattern. The ρ statistic quantifies the proportion of observed eigenvalues falling below the lower bound of the null distribution. See *SI Appendix* Fig. S2 for results on correlation matrices. (**B**) Visualization of the variation in matrix space based on EigenTensor Decomposition (ETD) analysis. Each point corresponds to a correlation matrix (100 for each of the 57 genera) projected onto the first two eigentensors (ETs), with vertical and horizontal bars representing the full range of observations for a given genus (100% credibility intervals). The percent variance explained by each ET is shown in parentheses on the axis. Cranium from MorphoSource (used with permission, see SI). High overlap in the matrix space indicates lower structural divergence among matrices. See *SI Appendix* Fig. S3 for results on covariance matrices. (**C**) Disparity-through-time (DTT) plot, illustrating how disparity in trait variance slowly accumulates across the primate phylogeny. 0 indicates no variation among clades, whereas 1 means all the disparity in trait covariance is partitioned among subclades. Matrix structure disparity (black lines) falls predominantly above the BM expectation, consistent with a pattern of evolutionary stasis. Despite deep-time divergence, all lineages share more similar covariance structures. Skull from MorphoSource, used with permission.

The 57 genus-level P-matrices were first compared using the Krzanowski common subspace method (Aguirre et al., 2014), which assesses the degree to which the leading subspaces of the matrices share a common orientation. The distribution of observed eigenvalues of **H** (λ) closely followed the null expectations across ranks, indicating substantial similarity in the orientation of the major covariance subspaces, broadly consistent with a common pattern across genera (Fig. 2A and *SI Appendix* Fig. S2). The low values of ρ further indicate that only a small fraction of λ fell below the lower bound of the null distribution, providing little evidence for reduced subspace alignment. However, this small fraction was concentrated primarily at the second rank of covariance matrices.

To further investigate the possible origins of these small departures detected in the Krzanowski method, we examined the shared matrix space using the EigenTensor Decomposition (ETD) analysis (Aguirre et al., 2014). ETD identifies the main axes of variation in matrix structure, thus providing a complementary view of how individual matrices are distributed within the broader space of covariance structures and revealing whether departures from a common structure are widespread or concentrated in a small number of taxa. Because ETD is scale-sensitive, we focused on correlation matrices rather than covariance matrices, allowing assessment of trait association patterns independent of trait variances. Our results showed a high overlap between matrices from all clades, with the majority of the variation in matrix structure explained by the first two tensors (ET1 = 60.79%, ET2 = 6.38%; Fig. 2B). Covariance matrices showed a similar result, but four genera overlapped slightly less with the remaining taxa (*Cebus, Nomascus, Symphalangus*, and *Pongo*; *SI Appendix* Fig. S3 for covariance matrix results). Together, these results suggest that the modest departures from common subspace alignment detected by the Krzanowski analysis are unlikely to reflect pervasive differences across the primate radiation and may instead be associated with variation in a relatively small subset of matrices.

We then performed a series of pairwise comparisons using the Random Skewers method (total of 1596 comparisons), which evaluates how similar two covariance matrices are with respect to their predicted evolutionary responses (Cheverud and Marroig, 2007). The high levels of similarity found among the comparisons (Table 1), not only within (RS median=0.868; sd=0.059) but also between major clades (RS median=0.791; sd=0.057), indicate minimal divergence in the covariance structure of all matrices (*SI Appendix* Fig. S4). An inspection of the distribution of variance for each covariance matrix indicates that primates follow the same trend: a higher proportion of variance is concentrated in the first eigenvector (mean=24.1%, sd=6.60), followed by a steady drop in the following directions (*SI Appendix* Fig. S5). We then tested whether traits grouped by developmental or functional hypotheses exhibit stronger covariation than would be expected by chance. This within- versus between-module comparison (AVG-ratio) provides a measure of the strength of modular organization in cranial trait covariation. The modularity analysis showed that all P-matrices exhibit similar (and higher than expected by chance) AVG-ratios for developmental and functional hypotheses, indicating a common modular structure among all genera (*SI Appendix* Fig. S6). The stasis in the covariance structure across all primates is also reinforced by a disparity-through-time (DTT) analysis on the eigentensor scores of the ETD (Fig. 2C). The DTT revealed a high, relatively flat curve, indicating limited differentiation in covariance structure among major lineages throughout the primate radiation. The empirical disparity curve remained above the BM expectation, a pattern consistent with evolutionary stasis rather than radiation.

**Table 1.** Primates show conserved evolutionary properties. Median and standard deviation of matrix similarity (Random Skewers) and three evolutionary potential metrics are summarized for the whole Order and by major taxonomic clade.

| Clade | matrix similarity (RS) | evolvability ($\log_{10}$) | autonomy | integration | number of matrices |
| --- | --- | --- | --- | --- | --- |
| Primates | $0.806 \pm 0.066$ | $-2.070 \pm 0.154$ | $0.271 \pm 0.030$ | $0.254 \pm 0.056$ | 57 |
| Strepsirrhini | $0.781 \pm 0.060$ | $-2.036 \pm 0.127$ | $0.275 \pm 0.034$ | $0.238 \pm 0.045$ | 14 |
| Platyrrhini | $0.804 \pm 0.061$ | $-2.153 \pm 0.110$ | $0.257 \pm 0.023$ | $0.250 \pm 0.050$ | 19 |
| Cercopithecidae | $0.813 \pm 0.066$ | $-2.068 \pm 0.138$ | $0.282 \pm 0.031$ | $0.280 \pm 0.065$ | 16 |
| Hominoidea | $0.793 \pm 0.061$ | $-1.935 \pm 0.192$ | $0.276 \pm 0.019$ | $0.236 \pm 0.048$ | 8 |

Interestingly, we found a positive correlation between shape divergence and matrix disparity, as measured through the ETD. However, this relationship is weak, and only a small fraction of the variation in matrix dissimilarity is explained by shape difference. Multiple regression on distance Matrices (MRM), including the phylogenetic distance matrix as a covariate (Legendre, Lapointe, and Casgrain, 1994), produced a regression coefficient of 0.139 (p< 0.001, Fig. 3; *SI Appendix* Fig. S7). These results indicate that mean shape divergence has evolved somewhat independently of divergence in matrix structure.

**Figure 3.**
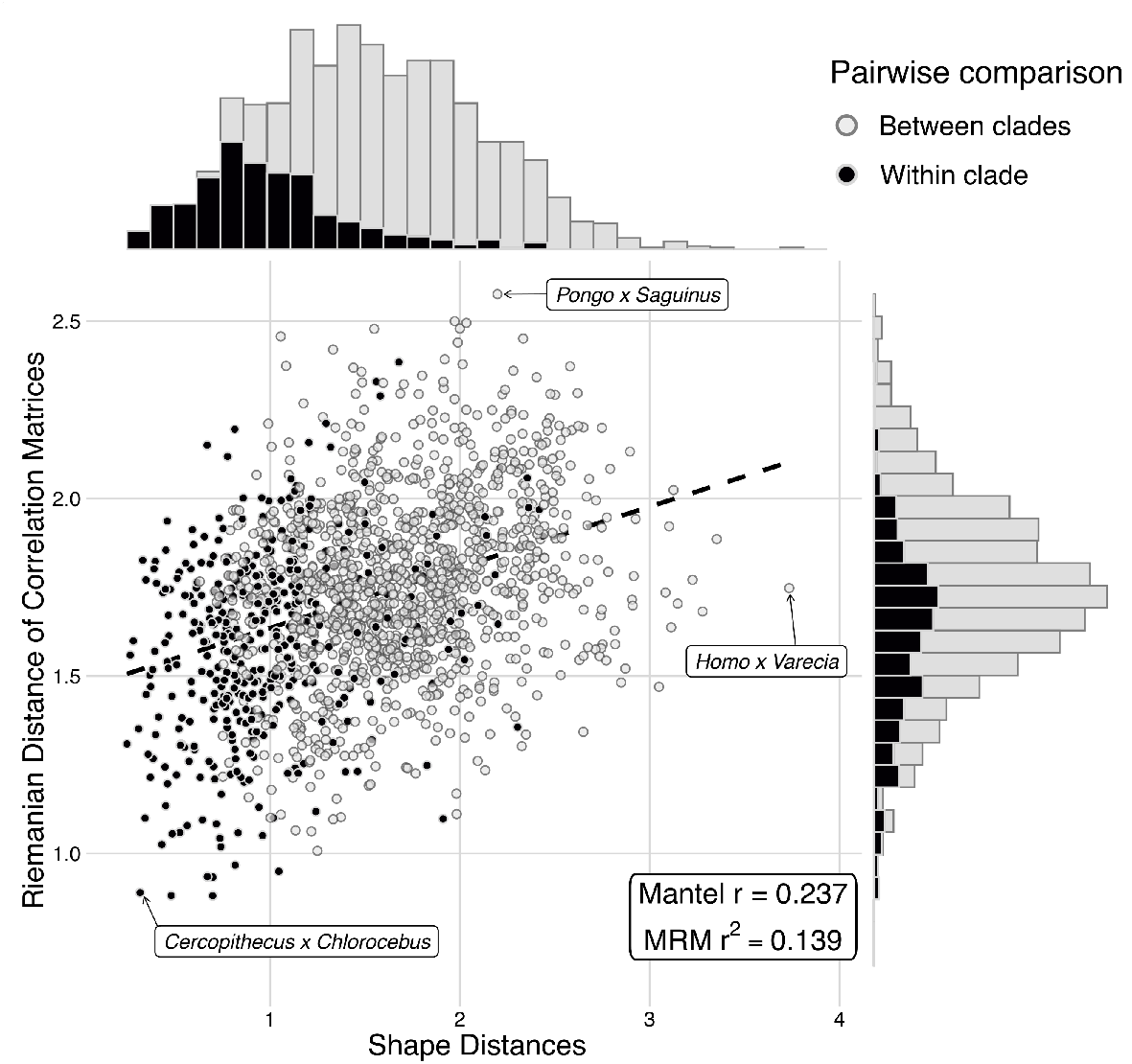
Matrix and shape divergence are weakly coupled. Each point represents the relationship between pairwise comparisons of mean matrix (Riemannian distance in the y-axis) and mean shape (Mahalanobis distance in the x-axis) of all primate genera (*N* = 57, for a total of 1596 comparisons). Density plots along the axes illustrate the distribution of distances within (black) and between (gray) major clades. Partial Mantel tests for dissimilarity accounting for the phylogenetic relationship (p=0.002, 999 permutations). Multiple regression on distance Matrices (MRM) using the phylogenetic distance matrix as a covariate. Labels with arrows point to extreme cases, but see *SI Appendix* Fig. S7 for all pairwise values.

### Similar evolutionary potential across primates

Using a simulation approach grounded in evolutionary quantitative genetics, we then compared the evolutionary properties of 57 primate genera by analyzing three evolutionary statistics derived from covariance matrices (Table 1; *SI Appendix* Fig. S8A): mean evolvability, autonomy, and integration (see methods for details). Mean evolvability represents the total genetic variation averaged across all morphospace directions, with the caveat that we are using the **P** matrices as a proxy for the **G** matrices (Holstad et al., 2024; Marroig and Cheverud, 2001a; Cheverud, 1988). Autonomy measures the proportion of genetic variation that is available to respond to directional selection, accounting for the influence of stabilizing selection (Hansen and Houle, 2008). Lastly, integration assesses how genetic variation is distributed across morphospace, with higher values indicating a concentration of variation in fewer directions, potentially due to constraints (Machado, Hubbe, et al., 2019; Marroig, Shirai, et al., 2009). Inspection of these matrix statistics reveals that the taxonomic groups largely overlap in their measures of evolvability, autonomy, and overall integration (Table 1; *SI Appendix* Fig S8A). The amount of phylogenetic signal was moderate for evolvability but negligible for autonomy and integration (*SI Appendix* Fig. S8B). Taken together with the results from ETD, RS, and DTT analyses, this suggests that at least some differences among P-matrices are attributable to the total amount of variation (evolvability) rather than to patterns of integration, modularity, and developmental constraints.

### Stability of evolvability despite pervasive directional selection

The remarkable similarity of cranial covariance and correlation patterns across primates raises a central question: how can such stability persist over the radiation of primates if directional selection is expected to reshape the phenotypic covariance at the microevolutionary scale (Turelli, 1988; Barnett, Meister, and Rainey, 2025; Jones, Arnold, and Bürger, 2007; Riedl, 1978)? One possible explanation is that directional selection is itself modular. Directional selection is thought to be able to reshape patterns of trait association through the differential fixation of pleiotropic alleles that reinforce the joint inheritance of selected traits (Melo and Marroig, 2015). If selection itself has a modular pattern that mirrors that of underlying trait associations, then selection would maintain, and possibly reinforce, these modular patterns (Melo and Marroig, 2015).

To test this hypothesis and investigate whether selection might have altered patterns of trait association, we reconstructed the distribution of multivariate selection gradients across the primate phylogeny. For that, we used a Bayesian implementation of phylogenetic independent contrasts (Martin and Weber, 2026). Under this framework, inferred selection gradients represent the amount of directional selection required to generate the observed phylogenetic divergence given the available covariance structure (Machado, 2020; Machado, Marroig, and Hubbe, 2022). This approach provides insight into what directions were preferentially selected throughout primate evolution (Machado, 2020; Simon, Machado, and Marroig, 2016). To summarize the information on past selection, we calculated the matrix of average cross-products of all selection gradients **Ω** (Machado, 2020), which is analogous to a covariance matrix of the realized selection gradients. Distribution of the eigenvalues of **Ω** showed that selection gradients were unevenly distributed across morphospace, indicating that some evolutionary directions were favored more frequently than others (Fig. 4A, *SI Appendix* Fig. S9). Trait loadings along the first two major axes of selection (analogous to Principal Components) revealed distinct patterns (Fig. 4B). The first axis indicates an increase in the neurocranial region along with generalized expansion of the facial region consistent with a pressure to increase in size, which is in line with previous findings for primates (Marroig and Cheverud, 2010; Marroig, Melo, and Garcia, 2012). The second axis captures variation associated with a contrast between the neurocranial and facial regions, consistent with variation in the cranial base flexion. These axes reflect the extreme variation in facial length and the increased level of encephalization observed in Primates (Lieberman, Ross, and Ravosa, 2000; Ross and Henneberg, 1995). We then asked whether certain directions of this selection gradient surface were more strongly associated than others. To examine this question, we tested if **Ω** had the same modular structure as our phenotypic matrices. Our results indicate strong support for modularity in the selective pattern, especially for the “functional” modular hypothesis (Fig. 4C), which proposes that cranial bones and sutures develop in response to the functional demands imposed by associated growing soft tissues, organs, cavities, and muscles (Moss and Young, 1960; Lieberman, 2011), but with weaker support for the developmental (neural crest/mesoderm) hypothesis. Taken together, these results suggest that correlated selection likely operated during the diversification of primate cranial traits, preferentially favoring trait combinations shaped by these functional-developmental processes, leading to a higher incidence of selection along certain directions of the morphospace that are predefined by cranial development.

**Figure 4.**
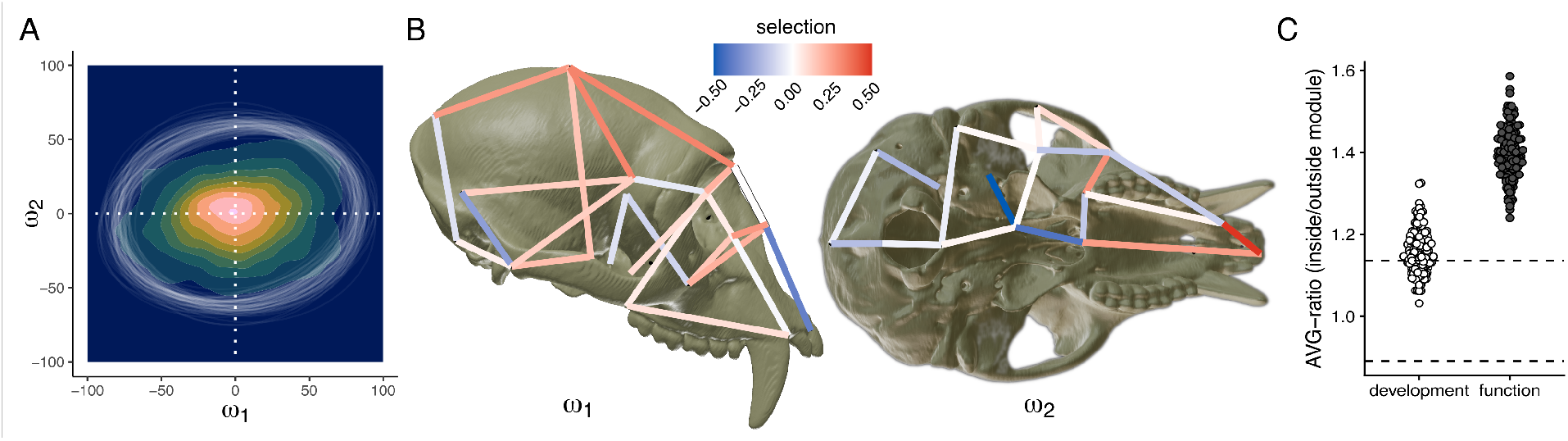
Directional selection is prevalent and aligned with cranial modular structure. (A) Kernel density of reconstructed branch-specific selection gradients projected onto the first two eigenvectors of the median realized selection surface. The covariance matrix of all selection gradients (**Ω**) is centered at the origin (white dashed lines) but slightly eccentric. Lighter pink indicates areas of higher density. White ellipses indicate the posterior distribution of **Ω** projected onto the median space. (B) Projection of the trait scores in the main axes of **Ω**. The first axis (in lateral view) shows an increase in encephalization and contrast between facial and neurocranial traits, and the second (ventral view) represents a flexion of the cranial base. Color gradient indicates the direction and strength of selection. (C) Modularity test on **Ω** showing that the cumulative multivariate directional selection that acted in the diversification of primates is highly modular and follows a similar organization as the skull’s developmental structure (*SI Appendix* Fig. S6). The dots represent the distribution of AVG ratios for the two modular hypotheses and the dashed lines indicate the interval expected for the null hypothesis of non-modular organization.

## Discussion

A central question in evolutionary biology is whether the genetic architecture that enables evolutionary change remains stable over prolonged time periods during which major radiations unfold. Here we found that primate cranial covariance structure (**P**-matrices) has remained remarkably conserved over tens of millions of years of evolution. The stability detected here does not imply that all these matrices are identical, but rather that their major directions of variation and their organization into functional and developmental modules have evolved within a relatively restricted realm of possibilities. Multiple complementary approaches converged on this result, including methods that compare matrix structure, evolvability, and the capacity of matrices to respond to selection. Because cranial **P**s and **G**s have been shown to be similar in primates (Marroig and Cheverud, 2010; Oliveira, Porto, and Marroig, 2009; Martínez-Abadías et al., 2009; Hubbe, Machado, et al., 2023; Roseman, Willmore, et al., 2010; Cheverud, 1996b), the similarity between **P**s across the entire radiation provides strong evidence that the underlying genetic architecture has also remained evolutionary stable (Marroig and Cheverud, 2001a; Hubbe, Melo, and Marroig, 2016). These findings challenge the widespread expectation in evolutionary quantitative genetics that additive genetic covariance structures (**G**-matrices) are too labile to retain predictive value over macroevolutionary timescales.

The persistence of a common covariance structure is striking because primates encompass one of the broadest ranges of cranial morphological diversification among mammals. Such diversification might be expected to alter patterns of trait covariation as directional selection can restructure covariances over microevolutionary timescales, specifically for the mammalian skull (Penna et al., 2017; Assis et al., 2017). Yet, our results suggest that primate diversification occurred largely through changes in mean cranial morphology rather than through repeated reorganization of the covariance structure. This distinction highlights an important aspect of evolvability: evolutionary potential depends on the distribution of heritable variation, not on the phenotype itself (Lande, 1979; Hansen, Houle, et al., 2023). Consequently, populations can undergo substantial morphological change while retaining a similar capacity to respond to selection, provided that genetic and phenotypic variation is preserved. Why then has this covariance structure remained so stable throughout primate evolution?

Two broad mechanisms could account for this pattern. The first is that covariance structure is itself maintained by internal stabilizing selection (Lande, 1976). Under this scenario, correlation among mutational effects (Jones, Arnold, and Bürger, 2007), pleiotropic effects with unmeasured traits (Bolstad et al., 2015), or the intricacies of developmental processes (Cheverud, 1996a) favor particular trait combinations. Given that other mammalian groups exhibit some degree of divergence in the integration pattern (Ferreira-Cardoso et al., 2022; Hubbe, Melo, and Marroig, 2016; Machado, Zahn, and Marroig, 2018; Machado, Hubbe, et al., 2019; Rossoni, Costa, et al., 2019), this would imply that primate skulls are under stronger internal selection, which seems unlikely. An alternative possibility is that the pattern of selection is preserved not because it’s actively constrained, but because directional selection repeatedly acts along the same developmental axes. If selection is aligned with the existing organization of variation, substantial morphological divergence could accumulate without requiring extensive reorganization of trait covariation.

Our results are more consistent with the second explanation. Our selection gradient reconstruction indicates that directional selection was highly structured during primate cranial evolution. The realized selection surface was distinctly eccentric, driven primarily by cranial base flexion and a contrast between the face and neurocranium (Fig. 4). Moreover, the modularity analysis of the realized selection gradient surface showed strong alignment with developmental processes related to the growth and function of the soft tissue and organs encased by the skull, suggesting that correlational selection favored certain trait combinations that act together during development (Sinervo and Svensson, 2002; Svensson et al., 2021; Moss and Young, 1960).

Thus, although directional selection has the potential to reshape patterns of integration in complex phenotypes (Melo and Marroig, 2015; Cai, Melo, and Des Marais, 2024), its tendency to align with pre-existing developmental axes of variation may limit its capacity to substantially disrupt primate cranial developmental trajectories. In contrast, other mammalian groups that showcase differences in integration, like canids and bats, seem to have experienced significant selection in directions not favored by integration patterns (Machado, 2020; Rossoni, Patterson, et al., 2024). Thus, while integration patterns can change over long timescales, selection must be sufficiently intense and discordant with those patterns to overcome the potential stabilizing force of internal selection. In primates, however, our results suggest that macroevolutionary change proceeds largely within persistent developmental constraints not only due to the effects of constraints (Rohner and Berger, 2023; Rohner and Berger, 2025), but because selection itself operates along the same directions. If selection itself repeatedly exploits the same developmental architecture, covariance structure is preserved because development and selection become aligned over macroevolutionary time.

More broadly, our results support extending quantitative-genetic theory to deep evolutionary timescales. Although these models were originally developed to describe evolutionary change within populations and have been applied successfully to divergence among recent lineages, particularly in studies of human evolution (Cramon-Taubadel, 2022; Schroeder and Ackermann, 2023; Ackermann, 2002; Roseman and Weaver, 2007; Roseman, 2004; Weaver, Roseman, and Stringer, 2008; Schroeder, Roseman, et al., 2014), their broader application has been limited by the assumption that **G**-matrices change too rapidly to retain predictive value over macroevolutionary scales. Increasing evidence has challenged this view by revealing continuity between micro- and macroevolution, particularly in the correspondence between within-population variation and rates or directions of divergence among lineages (Tsuboi et al., 2024; Houle et al., 2017; Rohner and Berger, 2023), a pattern also documented in primates (Marroig and Cheverud, 2004; Schroeder, Elton, and Ackermann, 2022). Our results extend this continuity one step further by suggesting that developmentally structured selection not only channels phenotypic diversification (Machado, Mongle, et al., 2023), but may also preserve the covariance architecture that determines future evolutionary responses. Quantitative-genetic models may therefore provide a mechanistic framework for understanding morphological evolution within few generations as well as trajectories of major evolutionary radiations.

## Materials and Methods

### Specimen sampling

We obtained morphometric measurements of 10,073 adult specimens, representing 309 out of 535 currently recognized species, 76 out of 84 genera, and all 16 Primate families for the most comprehensive taxonomic coverage of primate cranial morphology to date (*SI Appendix* Table S1).

### Cranial traits

To quantify primate cranial morphology, we used a 3D digitizer to record the 3D coordinates of 21 homologous landmarks defined at the intersection of bone sutures and other easily identifiable cranial structures (*SI Appendix* Table S2, and Fig. S10; Cheverud, 1984; Marroig and Cheverud, 2001b; Porto et al., 2009). We then calculated 37 Euclidean distances between landmark pairs (*SI Appendix* Table S3). These interlandmark distances capture general dimensions of singular bones and structures and have been broadly applied in previous investigations across mammals, including humans and nonhuman primates (Cheverud, 1996b; Marroig and Cheverud, 2001a; Oliveira, Porto, and Marroig, 2009; Porto et al., 2009; Schroeder, Elton, and Ackermann, 2022). The morphological variation in cranial shape and size was visually inspected using Principal Component Analysis (PCA) on the log-transformed data from the complete dataset (specimens of all species regardless of sample size). An isometric-size-free PCA was also performed using log-shape ratios defined as the logarithm of the ratio between each variable and the geometric mean of each individual.

### Phenotypic (co)variance and correlation matrices

To ensure robust estimation of trait variances and covariances, we calculated phenotypic covariance matrices (**P**) and their corresponding correlation matrices at the genus level, taking advantage of larger sample sizes available at this taxonomic scale (Grabowski and Porto, 2017). While P-matrices are not expected to be identical across species, empirical studies show that matrix similarity increases with phylogenetic relatedness (Marroig and Cheverud, 2001a; Machado, Zahn, and Marroig, 2018). Although covariance structures may vary due to ecological, developmental, or stochastic factors, pooling within-group variation across closely related species or populations provides a more reliable estimate of the underlying covariance architecture shaped by shared developmental and genetic constraints (Turelli, 1988). For genera with at least 39 specimens, we calculated a pooled within-genus **P**-matrix (**W**) by fitting a multivariate linear model with species as a fixed effect, for a total of 57 genus-level Ps. All Tarsiiform genera were below this threshold, so they were excluded from the P estimation. Although sexual dimorphism contributes to intraspecific variation, this investigation does not focus on its role in shaping covariance structure. Therefore, we excluded individuals with no sex information (N=213) and removed the confounding influence of sexual dimorphism by including sex as a fixed effect term in the linear model for Hominoidea, Cercopithecidae, and Platyrrhini clades. To account for uncertainty in the estimation of trait variances and covariances, we employed a Bayesian approach with a weakly informative regularized Wishart prior with a diagonal prior covariance matrix of the observed variances and number of traits plus one degrees of freedom (Melo, Garcia, et al., 2016). This Bayesian estimator provides regularization and many of the same advantages as a shrinkage estimator (Schäfer and Strimmer, 2005), while avoiding the issues that a naive maximum likelihood covariance matrix can cause when estimating selection gradients (Marroig, Melo, and Garcia, 2012). For each genus, we sampled 100 matrices from the posterior distribution, generating a distribution of possible covariance structures rather than relying solely on a point estimate of **P**. We assessed the self-similarity of each posterior sample by calculating the Principal Component Similarity (Melo, Garcia, et al., 2016) among each sample of the posterior and the median matrix (*SI Appendix* Tables S4-5).

All matrices were then mean-scaled by dividing each matrix by the outer product of the trait means obtained from the linear models (Hansen and Houle, 2008). This standardization allows for comparisons of covariation structure independent of absolute trait size, facilitating interpretation in terms of proportional variation, a necessary step when comparing covariance matrices across taxa that differ in overall size (Fig. 1B). We calculated correlation matrices from **P***S* by dividing each matrix by the outer product of the trait’s standard deviations. All analyses described below were conducted on both covariance and correlation matrices.

### Shared subspace comparison

We evaluated the similarity in the orientation of phenotypic (co)variance and correlation matrices across genera using the Krzanowski shared subspace comparison method (Aguirre et al., 2014; Melo, Garcia, et al., 2016). Briefly, this method tests the null hypothesis that all matrices under scrutiny share a similar structure by comparing the alignment of the subspaces spanned by their first leading eigenvectors. For each posterior sample of the genus-specific **P**s, we calculated the **H** matrix

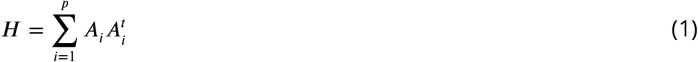

where **A**_*i*_ is a column matrix containing the leading eigenvectors of the *i*-th matrix being compared, *p* is the number of matrices being compared, and *t* denotes matrix transposition. The eigenvalues of **H** (λ) quantify the degree of shared orientation among the leading subspaces across matrices, with larger values indicating greater shared subspace orientation and the maximum value equals *p*, corresponding to complete overlap. To assess whether the observed degree of subspace alignment is exchangeable among genera, we generated a null distribution by bootstrapping residuals from the full dataset and re-estimating the posterior distributions of covariance matrices on randomized data (Aguirre et al., 2014). This procedure was repeated across 1,000 replicates, and the observed **H** matrix eigenvalues were then compared to this null distribution. For visualization, we plotted the posterior distribution of observed λ against a null envelope derived from randomized data for the first 15 ranks (Aguirre et al., 2014). The null envelope was calculated using rank-specific quantiles adjusted for multiple-comparisons, with a significance level α = 0.05/(2 ∗ *ranks*). Observed eigenvalues falling within the null envelope indicate high alignment in the subspaces that is compatible with the null hypothesis of a shared subspace, whereas values below the lower bound indicate less alignment than expected under randomization. We summarized the overall departure from the null by calculating ρ, the mean proportion of observed rank-specific λ falling below the lower bound of the null distribution. Thus, lower ρ values indicate that relatively few dimensions exhibit less subspace alignment than expected under randomization, suggesting high similarity in matrix orientation across taxa.

### Matrix Space

To simultaneously compare the pattern of variation in all **P** matrices, we constructed a matrix space using an Eigen Tensor Decomposition (ETD) analysis (Hine et al., 2009; Basser and Pajevic, 2007; Melo, Garcia, et al., 2016). Briefly, this multivariate approach decomposes the (co)variance structure of a set of matrices and identifies independent directions (eigentensors, ET) in the matrix space that describe the largest sources of variability in the data, each one describing a unique pattern of covariance between traits. Importantly, these independent eigentensors are biologically meaningful and can be interpreted as axes of differentiation representing selective pressures (Hine et al., 2009). As in PCA, the greater the variation associated with a given ET (calculated as a percentage of the sum of all eigenvalues), the more relevant that direction is for explaining variability in the matrices. To reduce data dimensionality and improve computational performance, we applied the ETD to the full set of posterior median (co)variance and correlation matrices projected onto a common basis as **P**_**r**_ = **V**^*t*^**PV**, where **P**_**r**_ is the rotated **P** into the common subspace and **V** are the eigenvectors of **W**, the pooled-within-group covariance matrix for the entire sample (Machado, Zahn, and Marroig, 2018). **W** was calculated using a linear model on the entire dataset, with sex and species as fixed effects and their interactions (*SI Appendix* Table S4 for a summary). To evaluate whether differences in cranial shape explain differences in the structure of trait covariation, we calculated the pairwise Riemannian distance (Mitteroecker and Bookstein, 2009) between covariance matrices on this matrix space and compared it to the Euclidean shape distance among genera, based on our size-free PCA.

### Disparity through time

To evaluate how patterns of variance and covariance among traits evolved, we applied a Disparity Through Time (DTT) analysis (Pennell et al., 2014) to the projections of posterior phenotypic covariance matrices in the EigenTensor Decomposition (ETD). This analysis allows us to verify whether disparity in the structure of trait covariance is concentrated between clades (subclade disparity approaches 0) or within clades (subclade disparity approaches 1). The posterior scores for the respective genera were randomly paired with a distribution of 100 posterior trees pruned from Wisniewski, Lloyd, and Slater (2022), yielding 100 datasets. For each dataset, we calculated the empirical disparity following standard practices (Pennell et al., 2014), except that instead of using a single tree to simulate the null expectations under a Brownian Motion (BM) model, here we performed a simulation for each posterior tree. We then interpolated the disparity values from the simulated datasets at evenly spaced time intervals, and computed a 95% confidence envelope (2.5th and 97.5th quantiles) across all trees. This allowed us to perform a DTT analysis accounting for both phylogenetic and measurement uncertainties. DTT values falling below the simulated null expectations are consistent with evolutionary radiation, while values above it are consistent with stasis.

### Evolutionary potential

In addition to comparing the geometric properties of the matrices, we applied quantitative genetics theory to assess their evolutionary potential (Hansen and Houle, 2008). Specifically, we simulated 1,000 random, normalized selection vectors (*β*) and computed two evolutionary statistics (evolvability and autonomy) based on the multivariate breeder’s equation (Δ**z** = **G***β*; where Δ**z** indicates the response to selection (Lande, 1979; Melo, Garcia, et al., 2016)). Evolvability (ē) was calculated for each *β* as the size of the projection of the covariance matrix onto the direction of selection

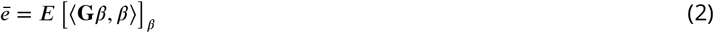

where ⟨.,. ⟩ represents the dot product between two vectors and *E* [.]_*β*_ indicates the expected value over random *β* vectors with unit norm. Then, the mean of all projections was taken as the overall evolutionary potential (i.e., average evolvability). This projection measures a population’s amount of additive genetic variance available in a particular direction of selection, averaged across all directions (Hansen and Houle, 2008). A related statistic is conditional evolvability, which measures the evolvability of a trait under the assumption of strong stabilizing selection on all other traits (Hansen and Houle, 2008). Because evolvability and conditional evolvability tend to scale together, we calculated the ratio between them, or “autonomy” (*ā*), as a measure of the proportion of the total variance on a trait that is able to evolve independently of other traits (Hansen and Houle, 2008). Analogously, the average autonomy across all random directions was used as an overall measure of relative trait independence.

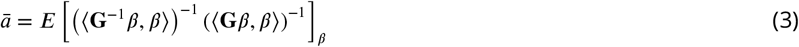

### Matrix pairwise similarity

We then compared each pair of covariance matrix using the Random Skewers (RS) method (Cheverud and Marroig, 2007), which consists of comparing two matrices with respect to their predicted evolutionary responses. For that, we employed the same simulation approach as described above to generate a thousand selection gradients (Melo, Garcia, et al., 2016), and then calculated the correlation between the respective responses to selection vectors when two matrices are subjected to the same selection gradients. The pairwise RS matrix similarity reported is the average cosine of the angle between the two Δ**z** vectors (or their vector correlation), computed using the same 1,000 normalized selection vectors (or skewers).

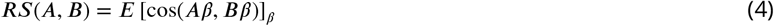

A value of 0 indicates that the two matrices are “completely unrelated,” whereas values close to 1 indicate that they behave similarly when subjected to the same selection gradient. Typically, values greater than 0.7 indicate a strong similarity in how matrices respond to selection.

### Phenotypic integration and modularity

Given that the structure of variance and the strength of trait correlations also influence the response to selection, we quantified the degree of trait integration and modularity. First, we computed the eigenvariance, a metric of phenotypic integration that quantifies how unevenly variance is distributed across principal components of the covariance matrix

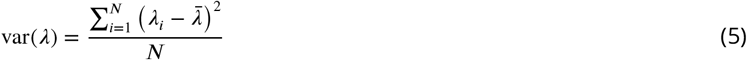

where λ are the eigenvalues of **P** and *N* are the number of traits. The final value was then scaled by the theoretical maximum integration possible for a matrix of similar rank (Machado, Hubbe, et al., 2019). Higher values indicate that variance is concentrated along fewer axes, consistent with stronger integration (Pavlicev, Cheverud, and Wagner, 2009). We calculated the eigen variance for 100 samples from the posterior distribution of each of the 57 P-matrices. Next, we evaluated whether functional or developmental integration could affect the co-inheritance and coordinated evolution of certain morphological traits by assessing whether correlations among trait groups exceeded those expected by chance. To that end, we tested two modular hypotheses that capture the relationship between traits arising from shared developmental origin (i.e., cranial neural crest or paraxial mesoderm) and from a shared functional-developmental mechanism (i.e., whereby bones respond to the growth of the enclosing soft tissue, organs, and cavities. *SI Appendix* Table S3, Fig. S6A) (Moss and Young, 1960; Cheverud, 1996a; Marroig, De Vivo, and Cheverud, 2004; Piekarski, Gross, and Hanken, 2014). Then, for each hypothesis, we calculated the ratio of average correlations within the same module (AVG+) to those outside the module (AVG-) using 100 samples from the posterior distribution of each P-matrix. To assess whether our empirical AVG ratio differed from chance, we performed the same operation, scrambling the rows and columns of the posterior matrices.

### Selection gradients

Under Lande’s multivariate breeder’s equation, evolutionary change (Δ**z**) reflects the joint effects of the structure of additive genetic (co)variance (**G**) and directional selection gradient *β*. Assuming that phenotypic covariance matrices (**P**) are good proxies for **G**, reconstructing evolutionary changes across the branches of the primate phylogeny allows estimation of the corresponding selection gradients. For that, we used a restructured version of Lande’s breeder’s equation (*β* = **G**^−1^Δ**z**). In macroevolutionary datasets, Δ**z** is not observed directly along every lineage, but we can approximate empirical estimates of evolutionary change using Phylogenetic Independent Contrasts (PICs) (Machado, 2020), which estimate the amount and direction of trait divergence accumulated between sister lineages after accounting for shared evolutionary history. To obtain these branch-specific changes, we implemented a Bayesian version of PICs by using Brownian Bridges to reconstruct a posterior distribution of ancestral states at each node (Martin and Weber, 2026) (see *SI Appendix* for details). We then summarized the covariance structure of all reconstructed selection gradients in a matrix (**Ω**) (Machado, 2020), which describes patterns of correlational selection. For visualization, we computed the leading eigenvectors of the median **Ω** that describe the main directions of selection. We also projected all *β*s onto this common base to produce a PCA-style visualization. We then tested the posterior distribution of **Ω**s for the same modularity hypotheses as for genus-level **P**s using the AVG-ratio and permutation tests.

## Data, Materials, and Software availability

The data and code used in this study will be made available for download via github prior to publication upon manuscript acceptance. For more details about the data, methodology, and results, please refer to the *Supporting Information Appendix*.

## Acknowledgments

This study relied on legacy data collected thanks to previous support from the Fundação de Amparo à Pesquisa do Estado de São Paulo (FAPESP; grants 2003/08706 − 8, 2004/13819 − 9, 2010/52469 − 4, 2011/14295 − 7, 2013/06577 − 8, and 2014/15116 − 7). AP was funded by the Sven och Lilly Lawskis fond för naturvetenskaplig forskning. A full list of museums housing the specimens used in this study is available in the Supporting Information. Alexandra Kralick provided an expert assessment of the flange status of male orangutans included in our sample. Lisandro Milocco, Erik Svensson, Masahito Tsuboi, and Ivan Prates provided thoughtful comments during the development of this project.

## Notes

### Competing Interest Statement

The authors have declared no competing interest.

