## Supporting_Information_Appendix for "Conserved covariance structure underlies 60 million years of morphological diversification in primates"

#### This PDF file includes:

Supporting text
Figs. S1 to S10
Tables S1 to S5
Legends for Dataset S1 to S2
SI References

#### Other supporting materials for this manuscript include the following:

Datasets S1 to S2

### Supporting Information Text

#### Materials.

**Specimen sampling.** We obtained morphometric measurements of 10,073 specimens, representing 309 out of 535 currently recognized species, 76 out of 84 genera, and all 16 Primate families for the most comprehensive taxonomic coverage of primate cranial morphology to date (Table S1). Our sample included only adult specimens to reduce potential bias in estimating covariances emerging from the grouping of different ontogenetic stages (1). Specimen age was assessed based on the complete fusion of basisphenoid and basioccipital sutures, full eruption of permanent canines and third molars, and overall morphology consistent with that of adults of each species. Taxonomic identifications were confirmed by crossing information on specimen sampling locality and the species' geographic ranges (2, 3); with nomenclature updates following (4).

This work was only possible thanks to the invaluable biological material safeguarded in 14 natural history collections of seven countries and three continents, as follows: *South-American Collections*: Museu Paraense Emilio Goeldi (**MPEG**, Belem, PA, Brazil), Museu de Zoologia da Universidade de Sao Paulo (**MZUSP**, Sao Paulo, SP, Brazil), Museu Nacional (**MZRJ**, Rio de Janeiro, RJ, Brazil); *North-American Collections*: American Museum of Natural History (**AMNH**, New York, NY, USA), Field Museum (**FMNH**, Chicago, IL, USA), Smithsonian National Museum (**USNM**, Washington, DC, USA), Harvard Museum of Comparative Zoology (**MCZ**, Cambridge, MA, USA), Duke Lemur Center Division of Fossil Primate Collection (**DFP**, Durham, NC, USA); *European Collections*: Naturalis Rijksmuseum van Natuurlijke Historie (**RMNH**, Leiden, Netherlands), Zoologisch Museum van Amsterdam (**ZMA**, Leiden, Netherlands), Muséum National d'Histoire Naturelle (**MNHN**, Paris, France), Royal Belgian Institute of Natural Sciences (**RBINS**, Brussels, Belgium), and Museum für Naturkunde (**ZMB**, **MfN**, Berlin, Germany), Natural History Museum, (**NHM**, London, UK).

#### Methods.

***3D Landmarks and Euclidean distances.*** To quantify primate cranial morphology, we captured the 3D coordinates for 21 cranial landmarks (Figure S10; Table S2) in adult specimens only. Landmarks were recorded using a Micoscribe MX (Immersion Corporation, San Jose, California) for Homo (by Arthur Porto (5)), Strepsirrhini and Taarsiiformes (by Anna Penna (6)); a Microscribe 3DX (Immersion Corporation, San Jose, California) for Catarrhini specimens (by Felipe Bandoni de Oliveira (7)); and a Polhemus 3Draw digitizer for Platyrrhini specimens (by Gabriel Marroig (8)).

We then calculated the Euclidean distances between selected pairs of landmarks for a total of 37 traits (see Table S3). Whenever the distance included at least one bilateral landmark (i.e., found on both sides of the cranium), we calculated the average between sides. These interlandmark distances capture general dimensions of singular bones and structures and have been broadly applied in previous quantitative genetics studies to describe the pattern of trait variance-covariance in mammals (5, 9–13). Moreover, these traits encompass a wide range of morphological features that have been extensively studied in the context of primate and Hominin cranial evolution (6, 7, 14–16). They include characteristics related to the size, shape, and positioning of the orbits; the length, width, and height of the face; the dimensions and configuration of the palate; the size and shape of the neurocranium; the width of the zygomatic arch and the degree of postorbital constriction; the form and orientation of the orbits and foramen magnum; as well as the flexion of the cranial base and the spatial arrangement between the neurocranium and splanchnocranium (17, 18). To ensure balanced sampling across taxa and maximize sample sizes, we measured intact or partially complete crania. In the few instances in which the area around a given bilateral landmark was damaged, we used the distances obtained from the intact side only.

**P-matrix estimation.** We assessed the self-similarity of each posterior sample pooled within-genus phenotypic matrix by calculating the Principal Component Similarity (19) among each sample of the posterior and the median matrix. For that, we projected each posterior sample onto the principal component space of a mean covariance matrix (pooled within-group matrix ($\mathbf{W}$, estimated from a global model including all specimens using the same Bayesian approach; see Table S4 for a summary) using eigenvector-based rotation, and calculated an adjusted similarity score by dividing the projected similarity by the self-similarity (Table S5).

### Selection gradients

Under Lande's multivariate breeder's equation (20), evolutionary change is determined jointly by the additive genetic covariance matrix $\mathbf{G}$ and the directional selection gradient $\beta$:

$$\Delta\mathbf{z} = \mathbf{G}\beta \qquad [S1]$$

Rearranging this relationship allows the expected selection gradient associated with an observed evolutionary change to be estimated as $\beta = \mathbf{G}^{-1}\Delta\mathbf{z}$. Because in macroevolutionary datasets $\Delta\mathbf{z}$ cannot be observed directly along extinct branches of the phylogeny, we approximated $\Delta\mathbf{z}$ by reconstructing branch-specific evolutionary changes derived from Phylogenetic Independent Contrasts (PICs). Following (21), these reconstructed changes were treated as empirical estimates of the multivariate response to selection accumulated along each branch.

First, we generated a standardized space for comparison by projecting all observations on the eigenvectors of the average within-group phenotypic covariance matrix $\mathbf{W}$ (15, 16, 21, 22). We then used species means and sampling variances in this space, along with a phylogenetic hypothesis of primate evolution (23), to estimate parameters for a multivariate evolutionary model using maximum likelihood (24). We fitted three models: a full multivariate BM model that jointly estimated trait rates of evolution and covariances; a simplified BM model that estimated only trait rates; and an Ornstein–Uhlenbeck process, which also accounted for stabilizing selection. Since the three models produced nearly identical rates of evolution for each eigenvector, we focus on the results from the simpler BM trait-rate-only model for clarity and interpretability. Parameter uncertainty was evaluated by fitting a BM model to 100 trees of the posterior sample. Uncertainty in phylogeny and model parameters was propagated by repeating the analyses across 100 trees sampled from the posterior distribution generated in (23).

Using these estimated evolutionary parameters, we reconstructed posterior distributions of ancestral phenotypes using stochastic character mapping based on Brownian bridges (25). This procedure simulates full probabilistic evolutionary histories for continuous traits that jointly incorporate phylogenetic uncertainty, uncertainty in evolutionary rates, and sampling error associated with species means. We generated a hundred simulations by combining the 100 posterior samples of the primate trees and their associated evolutionary rates, and the 100 posterior samples of the species-level covariance matrices. For species that didn't have enough samples for an adequate covariance matrix estimation, we assumed the maximum amount of measurement error by using the pooled-within-sample covariance matrix without scaling.

Rather than analyzing the complete simulated trajectories, we extracted the reconstructed phenotypic values at every node and calculated standardized PICs using Felsenstein's algorithm (26). These contrasts represent branch-specific estimates of multivariate evolutionary change after accounting for shared ancestry, that also take into account uncertainty in phylogeny, rates of evolution, and intraspecific errors. Using the distribution of PICs as an approximation for $\Delta\mathbf{z}$, we then used a restructured version of Lande's breeder's equation to estimate the corresponding selection gradients

$$\beta_i = (\mathbf{HWH})^{-1}\Delta\mathbf{z}_i \qquad \text{[S2]}$$

where $\mathbf{H}$ is a diagonal matrix containing the square roots of trait heritabilities $h^2$, and $\Delta z_i$ is the reconstructed evolutionary change associated with branch $i$. Following Cheverud's Conjecture (27), we assumed that i) phenotypic correlations provide reasonable approximations of the genetic correlations, allowing the within-species phenotypic covariance matrices to serve as a proxy for $\mathbf{G}$; and ii) that the covariance structure is stable across the group, which is supported by our results. To account for uncertainty in heritability, each posterior sample drew trait-specific $h^2$ from a uniform distribution between 0.3 and 0.6, encompassing the range typically reported for mammalian cranial traits (28–30).

Importantly, this procedure estimates one inferred selection gradient for each reconstructed branch of the phylogeny. Consequently, the resulting $\beta$ vectors represent the selection gradients that would be required to generate the reconstructed evolutionary changes under the assumed covariance structure. They should therefore be interpreted as branch-specific estimates of historical selection rather than direct observations of selection or complete temporal trajectories within individual evolutionary lineages.

To summarize the distribution of reconstructed selection gradients, we calculated the covariance matrix of all $\beta$ following (21):

$$\mathbf{\Omega} = \beta^t\beta\frac{1}{n-1} \qquad \text{[S3]}$$

Eigenanalysis of $\Omega$ identifies the principal directions in which inferred selection varied throughout primate evolution. The eigenvectors were subsequently projected back into trait space to facilitate biological interpretation.

To determine whether the inferred distribution of selection gradients departed from expectations under randomly oriented selection, we compared the eigenspectrum of the empirical $\Omega$ matrix with a null distribution generated from isotropic selection gradients. For the null model, we simulated multivariate selection gradients from a spherical multivariate normal distribution, ensuring that every direction in trait space was equally likely. These simulated gradients were converted into evolutionary responses using the average phenotypic covariance matrix and subsequently re-estimated using exactly the same analytical pipeline applied to the empirical data, including covariance estimation and reconstruction of selection gradients. This procedure generates the expected distribution of $\Omega$ eigenvalues under isotropic selection while accounting for uncertainty introduced by the estimation procedure itself. To evaluate the robustness of the inferred eigenspace, we projected every posterior sample of $\Omega$ onto the eigenvectors of the posterior median $\Omega$ and evaluated their overlap with zero.

**Table S1. Sample size for each primate genus considered in this study. Only genera with sample sizes larger than 39 were used to estimate phenotypic matrices. All species data were used to estimate selection gradients.**

| *Genus* | Clade | n species | sample size | avg specimens per species |
|---|---|---|---|---|
| *Arctocebus* | Strepsirrhini | 1 | 9 | 9.00 |
| *Perodicticus* | Strepsirrhini | 1 | 61 | 61.00 |
| *Loris* | Strepsirrhini | 2 | 26 | 13.00 |
| *Nycticebus* | Strepsirrhini | 4 | 41 | 10.25 |
| *Euoticus* | Strepsirrhini | 1 | 15 | 15.00 |
| *Otolemur* | Strepsirrhini | 1 | 59 | 59.00 |
| *Galago* | Strepsirrhini | 1 | 59 | 59.00 |
| *Daubentonia* | Strepsirrhini | 1 | 30 | 30.00 |
| *Varecia* | Strepsirrhini | 2 | 80 | 40.00 |
| *Lemur* | Strepsirrhini | 1 | 56 | 56.00 |
| *Prolemur* | Strepsirrhini | 1 | 16 | 16.00 |
| *Hapalemur* | Strepsirrhini | 4 | 70 | 17.50 |
| *Eulemur* | Strepsirrhini | 12 | 460 | 38.33 |

*Continues on next page*

*Continuation of Table S1*

| *Genus* | Clade | n species | sample size | avg specimens per species |
|---|---|---|---|---|
| *Indri* | Strepsirrhini | 1 | 66 | 66.00 |
| *Avahi* | Strepsirrhini | 3 | 64 | 21.33 |
| *Propithecus* | Strepsirrhini | 7 | 205 | 29.29 |
| *Lepilemur* | Strepsirrhini | 7 | 129 | 18.43 |
| *Phaner* | Strepsirrhini | 2 | 21 | 10.50 |
| *Cheirogaleus* | Strepsirrhini | 3 | 78 | 26.00 |
| *Allocebus* | Strepsirrhini | 1 | 1 | 1.00 |
| *Mirza* | Strepsirrhini | 1 | 21 | 21.00 |
| *Microcebus* | Strepsirrhini | 5 | 149 | 29.80 |
| *Cephalopachus* | Tarsiiformes | 1 | 18 | 18.00 |
| *Carlito* | Tarsiiformes | 1 | 18 | 18.00 |
| *Tarsius* | Tarsiiformes | 1 | 5 | 5.00 |
| *Pithecia* | Platyrrhini | 6 | 230 | 38.33 |
| *Cacajao* | Platyrrhini | 2 | 78 | 39.00 |
| *Chiropotes* | Platyrrhini | 4 | 153 | 38.25 |
| *Cheracebus* | Platyrrhini | 3 | 44 | 14.67 |
| *Callicebus* | Platyrrhini | 2 | 41 | 20.50 |
| *Plecturocebus* | Platyrrhini | 9 | 300 | 33.33 |
| *Alouatta* | Platyrrhini | 8 | 384 | 48.00 |
| *Brachyteles* | Platyrrhini | 2 | 40 | 20.00 |
| *Lagothrix* | Platyrrhini | 5 | 92 | 18.40 |
| *Ateles* | Platyrrhini | 4 | 226 | 56.50 |
| *Saimiri* | Platyrrhini | 6 | 257 | 42.83 |
| *Sapajus* | Platyrrhini | 5 | 325 | 65.00 |
| *Cebus* | Platyrrhini | 3 | 68 | 22.67 |
| *Aotus* | Platyrrhini | 7 | 179 | 25.57 |
| *Leontopithecus* | Platyrrhini | 2 | 37 | 18.50 |
| *Callimico* | Platyrrhini | 1 | 26 | 26.00 |
| *Callithrix* | Platyrrhini | 5 | 237 | 47.40 |
| *Cebuella* | Platyrrhini | 1 | 51 | 51.00 |
| *Mico* | Platyrrhini | 6 | 141 | 23.50 |
| *Leontocebus* | Platyrrhini | 8 | 153 | 19.12 |
| *Saguinus* | Platyrrhini | 11 | 399 | 36.27 |
| *Nomascus* | Hominoidea | 4 | 47 | 11.75 |
| *Hoolock* | Hominoidea | 1 | 47 | 47.00 |
| *Symphalangus* | Hominoidea | 1 | 47 | 47.00 |
| *Hylobates* | Hominoidea | 7 | 203 | 29.00 |
| *Pongo* | Hominoidea | 1 | 53 | 53.00 |
| *Gorilla* | Hominoidea | 2 | 233 | 116.50 |
| *Pan* | Hominoidea | 2 | 144 | 72.00 |
| *Homo* | Hominoidea | 1 | 237 | 237.00 |
| *Colobus* | Cercopithecidea | 5 | 317 | 63.40 |
| *Procolobus* | Cercopithecidea | 1 | 100 | 100.00 |
| *Piliocolobus* | Cercopithecidea | 10 | 365 | 36.50 |
| *Presbytis* | Cercopithecidea | 12 | 264 | 22.00 |
| *Simias* | Cercopithecidea | 1 | 29 | 29.00 |
| *Nasalis* | Cercopithecidea | 1 | 39 | 39.00 |
| *Pygathrix* | Cercopithecidea | 2 | 44 | 22.00 |
| *Rhinopithecus* | Cercopithecidea | 2 | 22 | 11.00 |
| *Semnopithecus* | Cercopithecidea | 8 | 109 | 13.62 |
| *Trachypithecus* | Cercopithecidea | 12 | 269 | 22.42 |
| *Allenopithecus* | Cercopithecidea | 1 | 3 | 3.00 |
| *Miopithecus* | Cercopithecidea | 2 | 53 | 26.50 |
| *Allochrocebus* | Cercopithecidea | 3 | 41 | 13.67 |
| *Erythrocebus* | Cercopithecidea | 1 | 34 | 34.00 |
| *Chlorocebus* | Cercopithecidea | 6 | 286 | 47.67 |
| *Cercopithecus* | Cercopithecidea | 20 | 656 | 32.80 |
| *Macaca* | Cercopithecidea | 19 | 604 | 31.79 |
| *Mandrillus* | Cercopithecidea | 2 | 45 | 22.50 |
| *Cercocebus* | Cercopithecidea | 5 | 112 | 22.40 |
| *Theropithecus* | Cercopithecidea | 1 | 33 | 33.00 |
| *Lophocebus* | Cercopithecidea | 2 | 53 | 26.50 |
| *Papio* | Cercopithecidea | 5 | 153 | 30.60 |

*End of Table S1*

**Table S2. Landmark descriptions and positions (* indicate landmarks in the midline). For a representation of the landmarks in a primate skull, see Fig. S10.**

| Landmark | Description | Position(s) |
|---|---|---|
| *IS | intradentale superior | midline |
| PM | premaxillary suture at the alveolus | right, left |
| *NSL | nasale | midline |
| *NA | nasion | midline |
| *BR | bregma | midline |
| PT | pterion | right, left |
| FM | fronto-malare | right, left |
| ZS | zygomaxillare superior | right, left |
| ZI | zygomaxillare inferior | right, left |
| MT | maxillary tuberosity | right, left |
| *PNS | posterior nasal spine | midline |
| APET | anterior petrous temporal | right and left |
| *BA | basion | midline |
| *OPI | opisthion | midline |
| EAM | anterior external auditory meatus | right, left |
| ZYGO | inferior zygo-temporal suture | right, left |
| TSP | temporo-spheno-parietal junction | right, left |
| TS | temporo-sphenoidal junction at the sphenoid | right, left |
| JP | jugular process | right, left |
| *LD | lambda | midline |
| AS | asterion | right, left |

**Table S3. List of distances used between pairs of landmarks (traits), and respective assignments to the substructure in functional and developmental modular hypotheses. The functional regions are related with the functional demands of the soft tissues encapsulated by the bones during development and through life. The developmental groups are subdivided into rostrofacial, corresponding to bones mostly derived from neural crest cells, and neurocranium, from the paraxial mesoderm. For more details see S6 A and (31–34).**

| Trait | Functional region | Developmental group |
|---|---|---|
| IS-PM | oral | rostrofacial |
| IS-NSL | nasal | rostrofacial |
| IS-PNS | oral, nasal | rostrofacial |
| PM-ZS | oral | rostrofacial |
| PM-ZI | oral | rostrofacial |
| PM-MT | oral | rostrofacial |
| NSL-NA | nasal | rostrofacial |
| NSL-ZS | nasal | rostrofacial |
| NSL-ZI | oral, nasal | rostrofacial |
| NA-BR | cranial vault | neurocranium |
| NA-FM | orbit | neurocranium |
| NA-PNS | nasal | rostrofacial |
| BR-PT | cranial vault | neurocranium |
| BR-APET | cranial vault | neurocranium |
| PT-FM | orbit | neurocranium |
| PT-APET | cranial vault | neurocranium |
| PT-BA | cranial vault | neurocranium |
| PT-EAM | zygomatic | rostrofacial |
| PT-ZYGO | zygomatic | rostrofacial |
| FM-ZS | orbit | neurocranium |
| FM-MT | zygomatic | rostrofacial |
| ZS-ZI | oral | rostrofacial |
| ZI-MT | oral | rostrofacial |
| ZI-ZYGO | zygomatic | rostrofacial |
| ZI-TSP | zygomatic | neurocranium |
| MT-PNS | oral | rostrofacial |
| PNS-APET | cranial base | neurocranium |
| APET-BA | cranial base | neurocranium |
| APET-TS | cranial base | neurocranium |
| BA-EAM | cranial base | neurocranium |
| ZYGO-TSP | zygomatic | rostrofacial |
| LD-AS | cranial vault | neurocranium |
| BR-LD | cranial vault | neurocranium |
| OPI-LD | cranial vault | neurocranium |
| PT-AS | cranial vault | neurocranium |
| JP-AS | cranial base | neurocranium |
| BA-OPI | cranial base | neurocranium |

**Table S4. First 5 principal components loadings for the pooled within-group covariance matrix ($W$).**

| | 1 | 2 | 3 | 4 | 5 |
|---|---|---|---|---|---|
| IS-PM | 0.186 | 0.051 | 0.021 | 0.057 | 0.082 |
| IS-NSL | 0.214 | 0.133 | 0.185 | 0.038 | 0.091 |
| IS-PNS | 0.192 | 0.050 | -0.015 | 0.056 | 0.086 |
| PM-ZS | 0.235 | 0.061 | -0.112 | -0.008 | 0.027 |
| PM-ZI | 0.183 | 0.024 | -0.078 | 0.071 | 0.262 |
| PM-MT | 0.134 | 0.014 | -0.009 | 0.036 | 0.075 |
| NSL-NA | 0.204 | -0.130 | -0.879 | -0.158 | -0.164 |
| NSL-ZS | 0.170 | 0.013 | -0.111 | -0.048 | -0.059 |
| NSL-ZI | 0.157 | -0.005 | -0.050 | 0.055 | 0.199 |
| NA-BR | 0.102 | -0.006 | 0.131 | 0.227 | -0.076 |
| NA-FM | 0.119 | -0.016 | 0.057 | 0.014 | 0.026 |
| NA-PNS | 0.157 | -0.008 | -0.054 | 0.037 | 0.040 |
| BR-PT | 0.049 | -0.052 | 0.004 | 0.213 | -0.014 |
| BR-APET | 0.083 | -0.044 | 0.040 | 0.094 | 0.031 |
| PT-FM | 0.224 | 0.759 | -0.007 | 0.074 | -0.063 |
| PT-APET | 0.123 | -0.244 | 0.116 | -0.042 | 0.003 |
| PT-BA | 0.134 | -0.176 | 0.087 | -0.025 | 0.016 |
| PT-EAM | 0.167 | -0.331 | 0.125 | -0.095 | 0.007 |
| PT-ZYGO | 0.232 | -0.268 | 0.167 | -0.148 | -0.069 |
| FM-ZS | 0.092 | -0.095 | -0.005 | -0.013 | -0.010 |
| FM-MT | 0.189 | 0.027 | 0.007 | 0.037 | 0.008 |
| ZS-ZI | 0.178 | -0.026 | -0.081 | 0.146 | 0.433 |
| ZI-MT | 0.176 | 0.024 | 0.022 | 0.102 | 0.067 |
| ZI-ZYGO | 0.226 | 0.041 | 0.154 | -0.192 | -0.547 |
| ZI-TSP | 0.216 | 0.033 | 0.083 | -0.128 | -0.284 |
| MT-PNS | 0.210 | 0.038 | 0.022 | 0.024 | 0.025 |
| PNS-APET | 0.165 | 0.066 | 0.058 | -0.126 | -0.127 |
| APET-BA | 0.167 | 0.015 | 0.014 | 0.029 | 0.018 |
| APET-TS | 0.129 | -0.074 | 0.009 | 0.063 | 0.060 |
| BA-EAM | 0.139 | -0.008 | 0.022 | 0.039 | 0.025 |
| ZYGO-TSP | 0.251 | -0.034 | 0.126 | -0.133 | -0.083 |
| LD-AS | 0.075 | -0.092 | -0.002 | 0.293 | -0.037 |
| BR-LD | 0.070 | 0.081 | 0.075 | -0.558 | 0.441 |
| OPI-LD | 0.104 | -0.150 | -0.027 | 0.510 | -0.086 |
| PT-AS | 0.122 | -0.190 | 0.064 | -0.052 | 0.060 |
| JP-AS | 0.133 | -0.044 | 0.014 | 0.056 | 0.067 |
| BA-OPI | 0.035 | -0.021 | 0.023 | -0.139 | 0.053 |
| **Proportion of Variance** | 0.277 | 0.068 | 0.064 | 0.054 | 0.049 |
| **Cumulative Proportion** | 0.277 | 0.345 | 0.409 | 0.463 | 0.512 |

**Table S5. comparison between matrices projected onto the first 15 axes of the global pooled within-group P matrix. *Self-similarity*– the average self-similarity between the median matrix and the distribution of matrices. *Similarity*– The raw similarity between the original and the projected matrices. *Similarity (adjusted)*– The adjusted similarity between the original and the projected matrices. Values were adjusted by the average self-similarity. Values between parentheses are the $95\%$ range of the highest density of the posterior.**

| *Genus* | Self-similarity | Similarity | Similarity (adjusted) |
|---|---|---|---|
| *Allochrocebus* | 0.867 | 0.879(0.828-0.908) | 1.000(0.956-1.000) |
| *Alouatta* | 0.992 | 0.978(0.974-0.981) | 0.986(0.981-0.989) |
| *Aotus* | 0.969 | 0.912(0.901-0.919) | 0.941(0.929-0.948) |
| *Ateles* | 0.977 | 0.948(0.939-0.955) | 0.970(0.960-0.977) |
| *Avahi* | 0.891 | 0.803(0.776-0.821) | 0.901(0.870-0.921) |
| *Brachyteles* | 0.834 | 0.841(0.797-0.881) | 1.000(0.956-1.000) |
| *Cacajao* | 0.933 | 0.915(0.887-0.932) | 0.980(0.951-0.999) |
| *Callicebus* | 0.868 | 0.833(0.782-0.861) | 0.960(0.901-0.992) |
| *Callithrix* | 0.975 | 0.927(0.918-0.934) | 0.951(0.941-0.958) |
| *Cebuella* | 0.881 | 0.812(0.783-0.840) | 0.921(0.889-0.953) |
| *Cebus* | 0.925 | 0.915(0.874-0.931) | 0.988(0.944-1.000) |
| *Cercocebus* | 0.962 | 0.950(0.934-0.962) | 0.988(0.971-1.000) |
| *Cercopithecus* | 0.995 | 0.979(0.977-0.981) | 0.984(0.982-0.986) |
| *Cheirogaleus* | 0.956 | 0.942(0.929-0.954) | 0.985(0.972-0.998) |
| *Cheracebus* | 0.855 | 0.829(0.771-0.856) | 0.969(0.901-1.000) |
| *Chiropotes* | 0.964 | 0.927(0.912-0.936) | 0.962(0.946-0.971) |
| *Chlorocebus* | 0.986 | 0.969(0.964-0.973) | 0.983(0.978-0.987) |
| *Colobus* | 0.985 | 0.958(0.952-0.962) | 0.972(0.966-0.976) |
| *Eulemur* | 0.990 | 0.926(0.923-0.930) | 0.936(0.932-0.939) |
| *Galago* | 0.941 | 0.868(0.842-0.886) | 0.923(0.895-0.941) |
| *Gorilla* | 0.979 | 0.904(0.894-0.910) | 0.923(0.913-0.930) |
| *Hapalemur* | 0.902 | 0.809(0.779-0.834) | 0.898(0.864-0.926) |
| *Homo* | 0.976 | 0.918(0.910-0.926) | 0.941(0.932-0.949) |
| *Hoolock* | 0.862 | 0.829(0.796-0.859) | 0.961(0.923-0.996) |
| *Hylobates* | 0.971 | 0.927(0.915-0.933) | 0.955(0.943-0.961) |
| *Indri* | 0.897 | 0.782(0.759-0.802) | 0.871(0.846-0.894) |
| *Lagothrix* | 0.937 | 0.918(0.891-0.933) | 0.980(0.951-0.995) |
| *Lemur* | 0.904 | 0.859(0.823-0.880) | 0.950(0.910-0.973) |
| *Leontocebus* | 0.960 | 0.907(0.896-0.916) | 0.945(0.934-0.954) |
| *Lepilemur* | 0.960 | 0.865(0.853-0.875) | 0.901(0.888-0.911) |
| *Lophocebus* | 0.932 | 0.907(0.879-0.920) | 0.973(0.943-0.987) |
| *Macaca* | 0.996 | 0.984(0.982-0.985) | 0.987(0.986-0.989) |
| *Mandrillus* | 0.898 | 0.825(0.787-0.847) | 0.919(0.877-0.943) |
| *Mico* | 0.951 | 0.846(0.831-0.861) | 0.889(0.874-0.906) |
| *Microcebus* | 0.951 | 0.838(0.819-0.855) | 0.881(0.861-0.899) |
| *Miopithecus* | 0.929 | 0.907(0.873-0.923) | 0.976(0.939-0.993) |
| *Nomascus* | 0.866 | 0.883(0.845-0.905) | 1.000(0.976-1.000) |
| *Nycticebus* | 0.870 | 0.824(0.772-0.845) | 0.947(0.887-0.971) |
| *Otolemur* | 0.938 | 0.876(0.852-0.890) | 0.934(0.908-0.949) |
| *Pan* | 0.959 | 0.893(0.878-0.904) | 0.931(0.915-0.943) |
| *Papio* | 0.986 | 0.932(0.927-0.936) | 0.945(0.940-0.949) |
| *Perodicticus* | 0.927 | 0.910(0.877-0.929) | 0.981(0.945-1.000) |
| *Piliocolobus* | 0.987 | 0.960(0.955-0.963) | 0.972(0.967-0.976) |
| *Pithecia* | 0.980 | 0.952(0.943-0.957) | 0.971(0.962-0.977) |
| *Plecturocebus* | 0.985 | 0.965(0.960-0.969) | 0.980(0.974-0.983) |
| *Pongo* | 0.949 | 0.914(0.895-0.928) | 0.963(0.943-0.978) |
| *Presbytis* | 0.971 | 0.932(0.925-0.938) | 0.960(0.952-0.966) |
| *Procolobus* | 0.956 | 0.916(0.900-0.926) | 0.958(0.941-0.968) |
| *Propithecus* | 0.973 | 0.904(0.894-0.910) | 0.929(0.918-0.935) |
| *Pygathrix* | 0.866 | 0.856(0.807-0.892) | 0.987(0.931-1.000) |
| *Saguinus* | 0.985 | 0.906(0.900-0.910) | 0.920(0.914-0.924) |
| *Saimiri* | 0.989 | 0.953(0.946-0.956) | 0.964(0.957-0.967) |
| *Sapajus* | 0.990 | 0.965(0.961-0.968) | 0.975(0.970-0.977) |
| *Semnopithecus* | 0.958 | 0.948(0.935-0.959) | 0.990(0.975-1.000) |
| *Symphalangus* | 0.877 | 0.892(0.846-0.918) | 1.000(0.965-1.000) |
| *Trachypithecus* | 0.978 | 0.940(0.933-0.944) | 0.961(0.954-0.965) |
| *Varecia* | 0.924 | 0.871(0.842-0.887) | 0.943(0.912-0.960) |

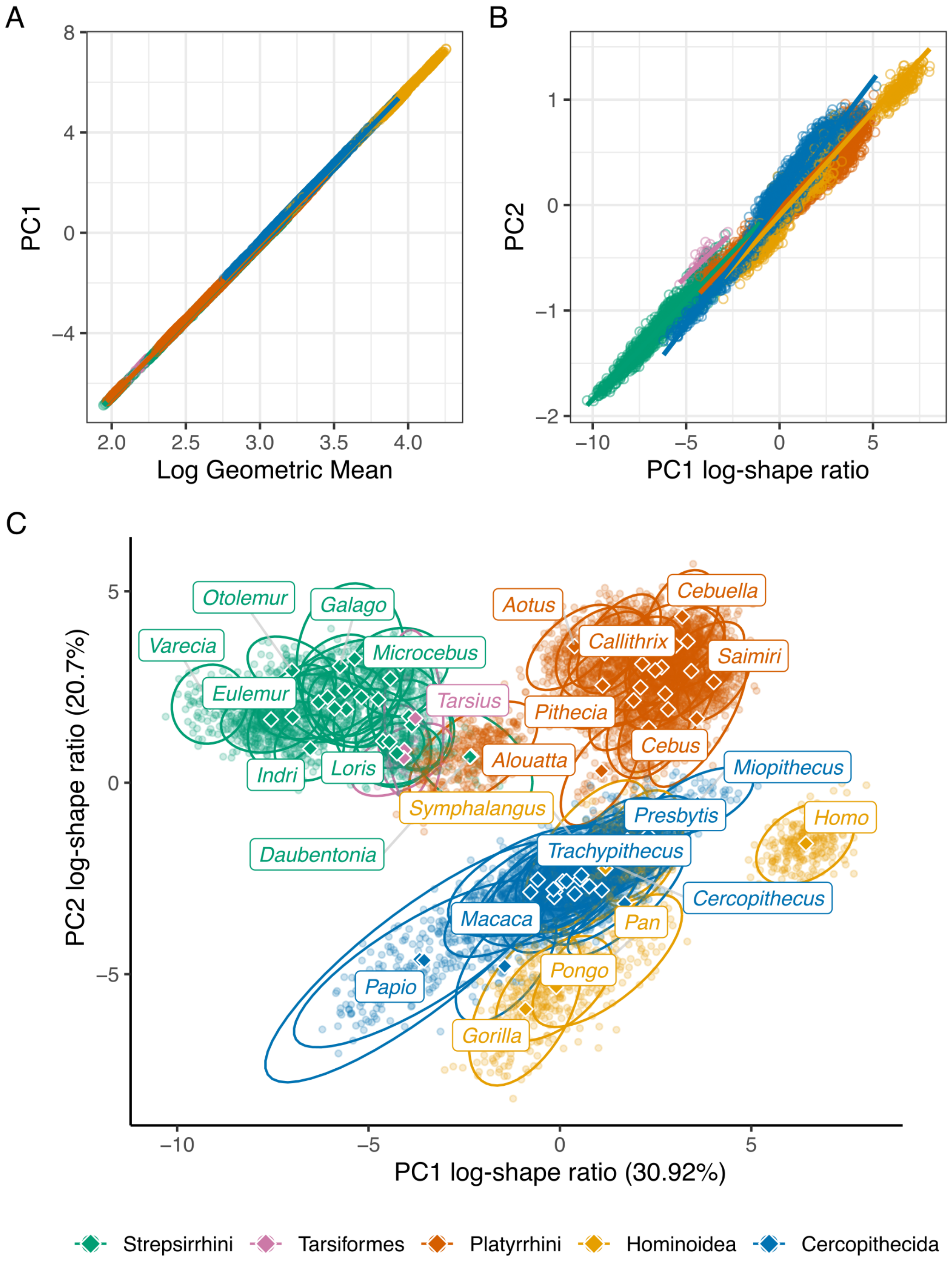


**Fig. S1.** (**A**) Regression of PC1 scores against the logarithm of the geometric mean of all cranial measurements. (**B**) Regression of PC2 scores against the PC1 scores of a PCA on isometric-size-free data using log-shape ratios defined as the logarithm of the ratio between each variable and the geometric mean of each individual. This transformation keeps all allometric-related shape variation. (**C**) Principal component analysis (PCA) on size-corrected data.

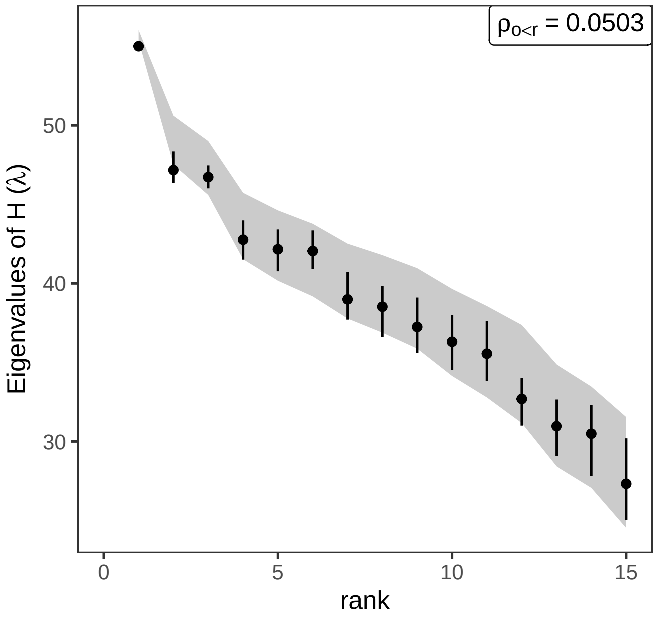


**Fig. S2.** Comparison of the 57 correlation matrices using the Krzanowski common subspace. Rank-specific eigenvalues of $\mathbf{H}$ $(\lambda)$ from the empirical posterior distribution (black) compared to the null envelope derived from randomized matrices (gray polygon). The dots represent posterior medians, and vertical bars indicate $95\%$ highest posterior density intervals. Eigenvalues approaching 57, the total number of matrices, indicate stronger shared subspace alignment across genera, whereas lower eigenvalues indicate weaker commonality among matrices. Overlap between empirical and null distributions indicates that the observed degree of subspace alignment is compatible with the randomization null and does not require genus-specific covariance structure to explain the observed pattern. The $\rho$ statistic quantifies the mean proportion of observed eigenvalues falling below the lower bound of the corresponding null distribution.

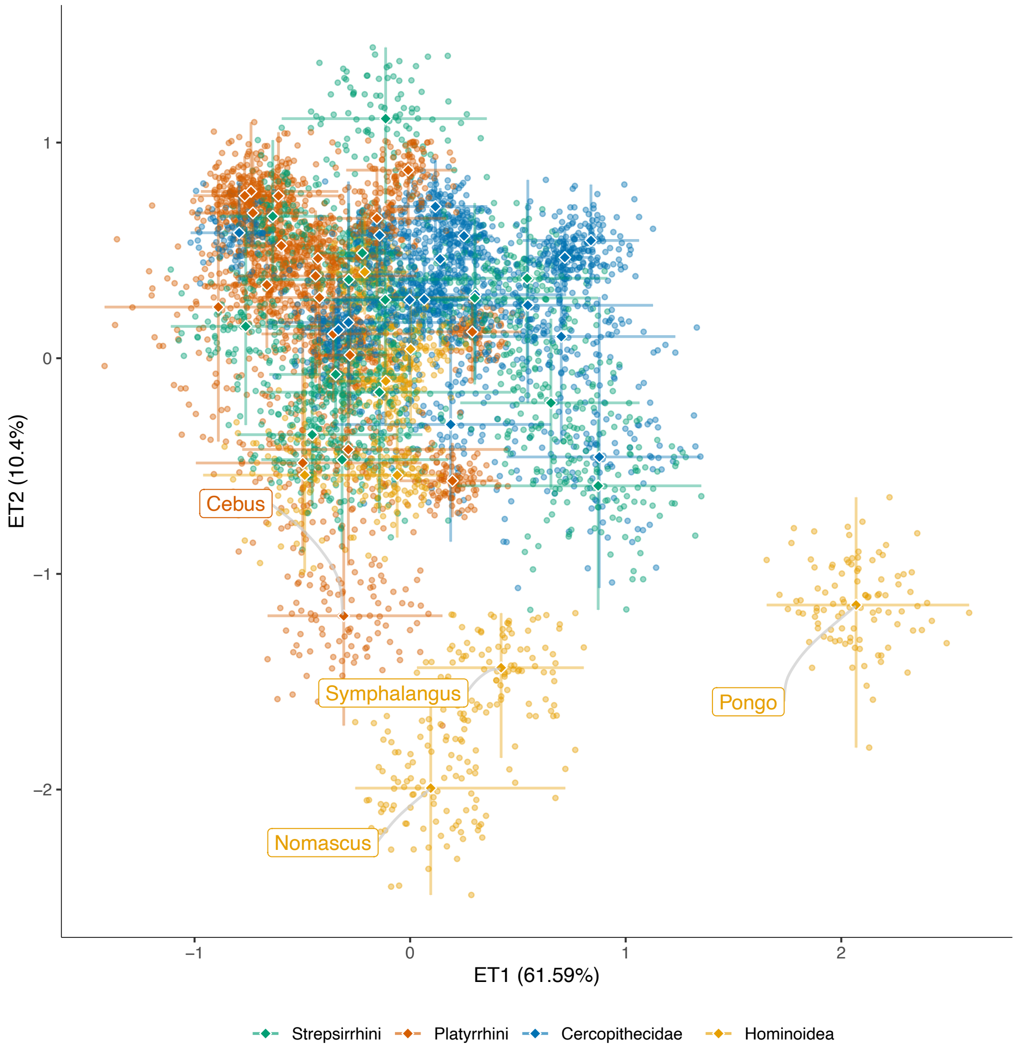


**Fig. S3.** Visualization of covariance matrix variation in matrix space based on EigenTensor Decomposition (ETD) analysis. Each point corresponds to a covariance matrix (100 matrices per genus, for a total of 5,700 points) projected onto the first two eigentensors (ET1 and ET2). To establish a common reference frame for comparison, we used the eigenvectors of $\mathbf{W}$ as a common-projection basis for all the posterior $(\mathbf{P}s)$ and median covariance matrices for all genera. Percent variance explained by each ET is indicated in parentheses. In our cranial dataset, the first eigenvector of the covariance matrices correspond to an allometric size axis.Therefore, different from an EDT analysis using correlation matrices (see main text Fig. 2B), here the first eigen tensor mostly captures among-taxon differences in size-related variance. Most genera cluster tightly in matrix space, consistent with the high similarity among correlation and covariance structures. A small number of genera occupy more isolated positions, likely reflecting differences in the proportion of variance associated with size of particular traits. For instance *Pongo*, is separated primarily along ET1, which may reflect its comparatively high proportion of variance associated with size ($35\%$; see Fig. S5), and higher evolvability, which reflects the total amount of variance (S8), not a change in the pattern of integration. One possible concern was that the disjunct distribution of *Pongo* in the ETD analysis, along with elevated size-related and total variance, could be due to uncontrolled ontogenetic effects in adult male morphotypes. While unflanged males differ substantially in body size and facial proportions from flanged males, their facial proportions are more similar to adult females (35). Not accounting for such differences in male shape and size could potentially bias our estimates. However, this explanation is unlikely in our dataset. Our sample contains a much higher proportion of confirmed flanged-to-unflanged males ($N = 36$ and $3$, respectively; Alexandra Kralick, personal communication), and males ($N = 39$) and females ($N = 14$) occupy distinct, non-overlapping regions of cranial morphospace (Supporting Dataset 1). Thus, the distinctive position of *Pongo* in matrix space is unlikely to result from ontogenetic variation in the expression of sexual dimorphism. More generally, it can be attributed to its higher total variance, as matrix eccentricity alone appears insufficient to explain matrix divergence in the ET analysis. Other genera, such as *Papio* and *Macaca*, showed even higher eccentricity, with larger proportions of variance associated with PC1 ($47\%$ and $41\%$, respectively), yet exhibited average levels of matrix similarity relative to the rest of the dataset. The remaining genera that sowed less overlap with other taxa in ET2 could reflect taxonomic uncertainty, for example, if individuals of *Symphalangus*, *Nomascus*, and *Cebus* were classified under outdated or inconsistent taxonomic schemes.

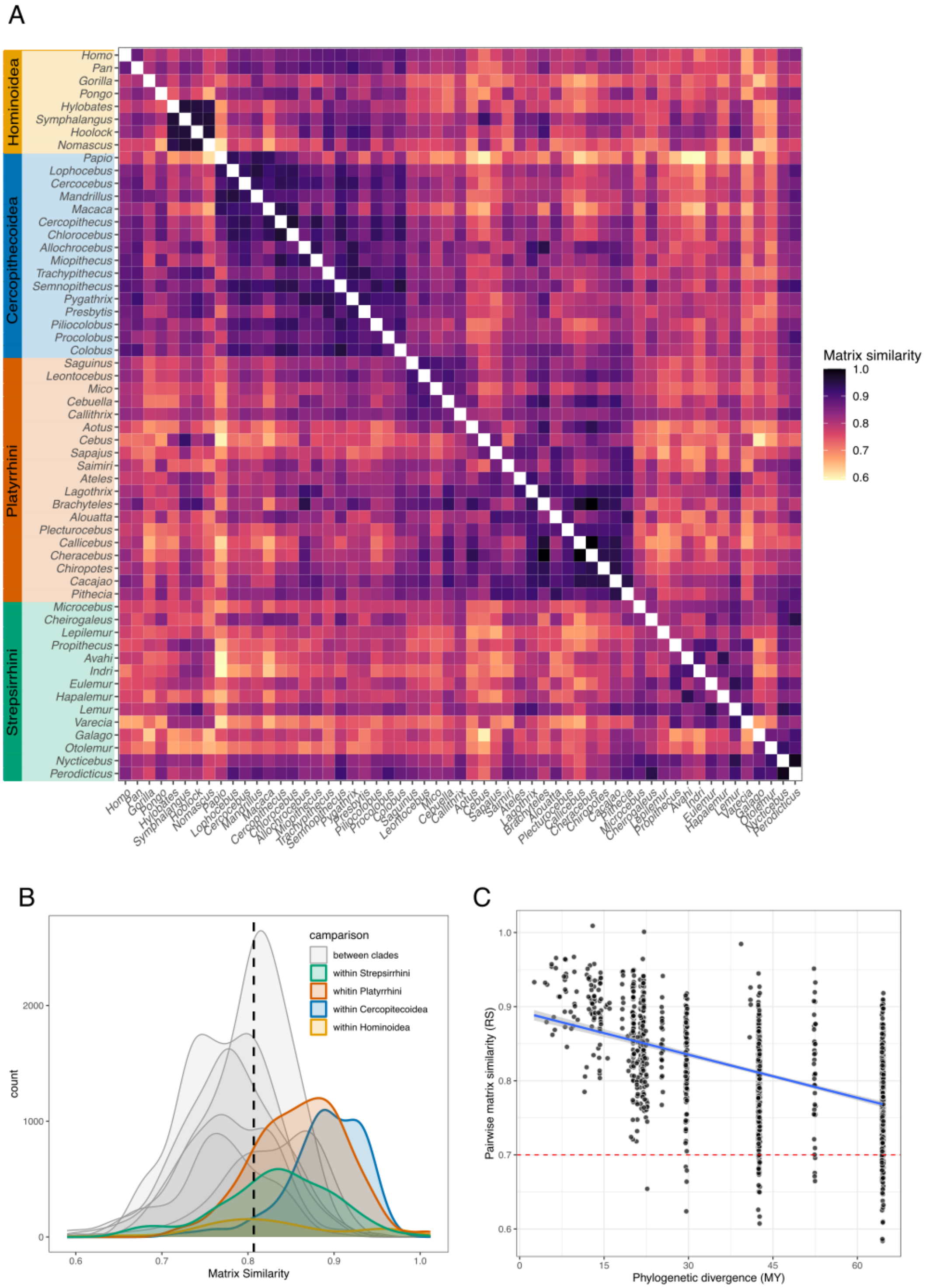


**Fig. S4.** Pairwise comparisons between the 57 covariance matrices. (**A**) Heath map showing the mean values of matrix similarity calculated using the Random Skewers method (total $= 1596$ comparisons). Darker colors indicate greater similarity, and colored bars to the left denote the major clades. (**B**) Distribution of pairwise comparison values between matrices from different clades (curves in gray) or within the same clade (colored following the same scale as the bar in panel A). The dashed line indicates the median pairwise matrix similarity for the complete dataset. (**C**) Pairwise similarity in cranial covariance structure declines significantly with increasing phylogenetic divergence (Mantel statistic r $= -0.544$; Pearson correlation $= -0.53$, $p < 0.01$). Typically, random skewers comparison values greater than 0.7 indicate strong similarity in how matrices respond to selection. Values below this threshold (only $5.45\%$ of comparisons) are restricted to comparisons among taxa that diverged more than 20 million years ago– yet, even among the deeply diverged taxa, similarity values remain higher than 0.7 for the majority of cases. This suggests that although closely related genera tend to have higher similarity, the structure of cranial phenotypic variance is remarkably stable across much of primate evolutionary history.

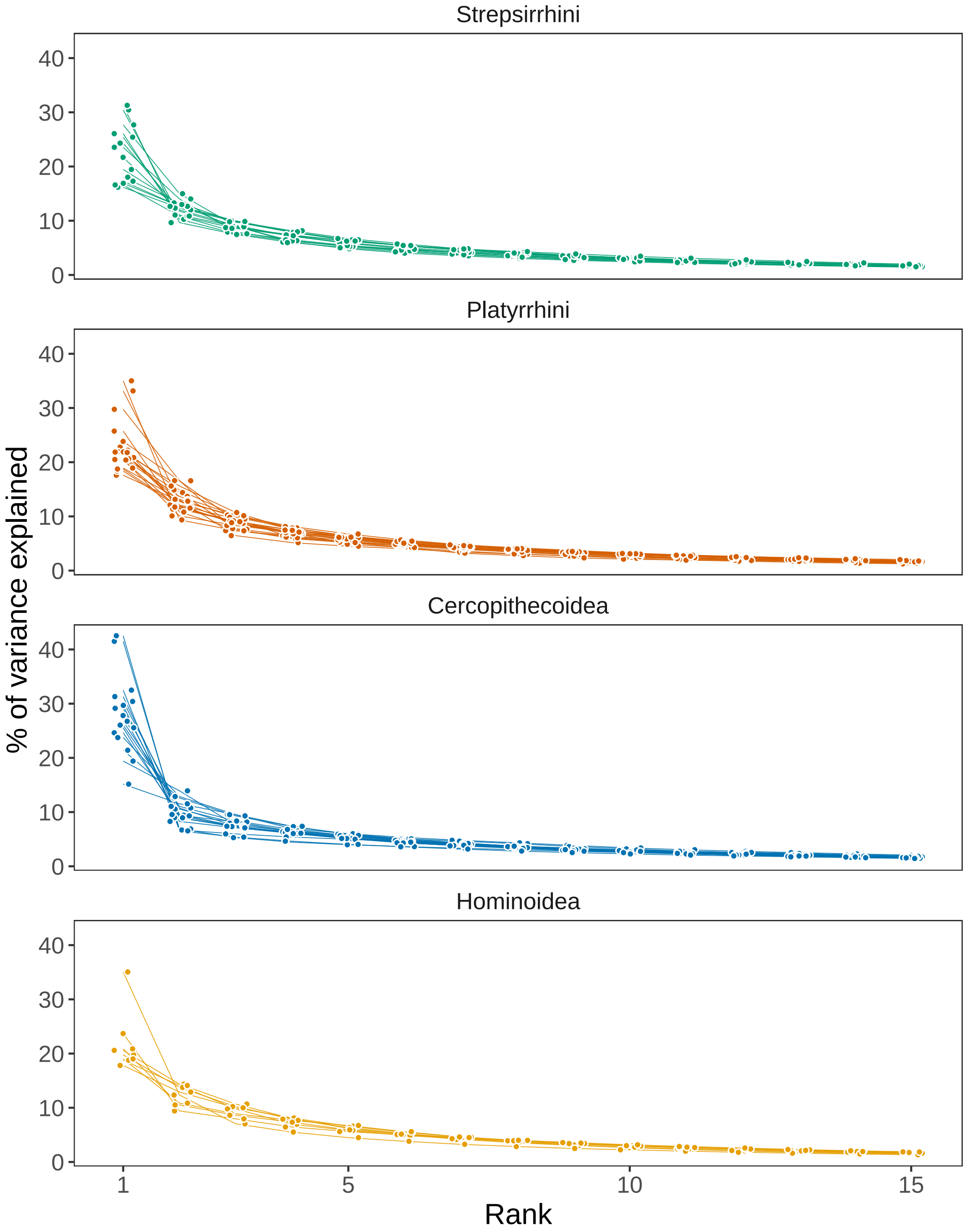


**Fig. S5.** Primates show a similar distribution of variance in each principal component. Although the first principal component accounts for most of the variance (mean=$24.1\%$, sd=$6.61$). Phenotypic integration values were strongly correlated with the proportion of variance explained by the first eigenvector (Pearson correlation=$0.961$, p$< 0.001$; $r^2 = 0.923$).

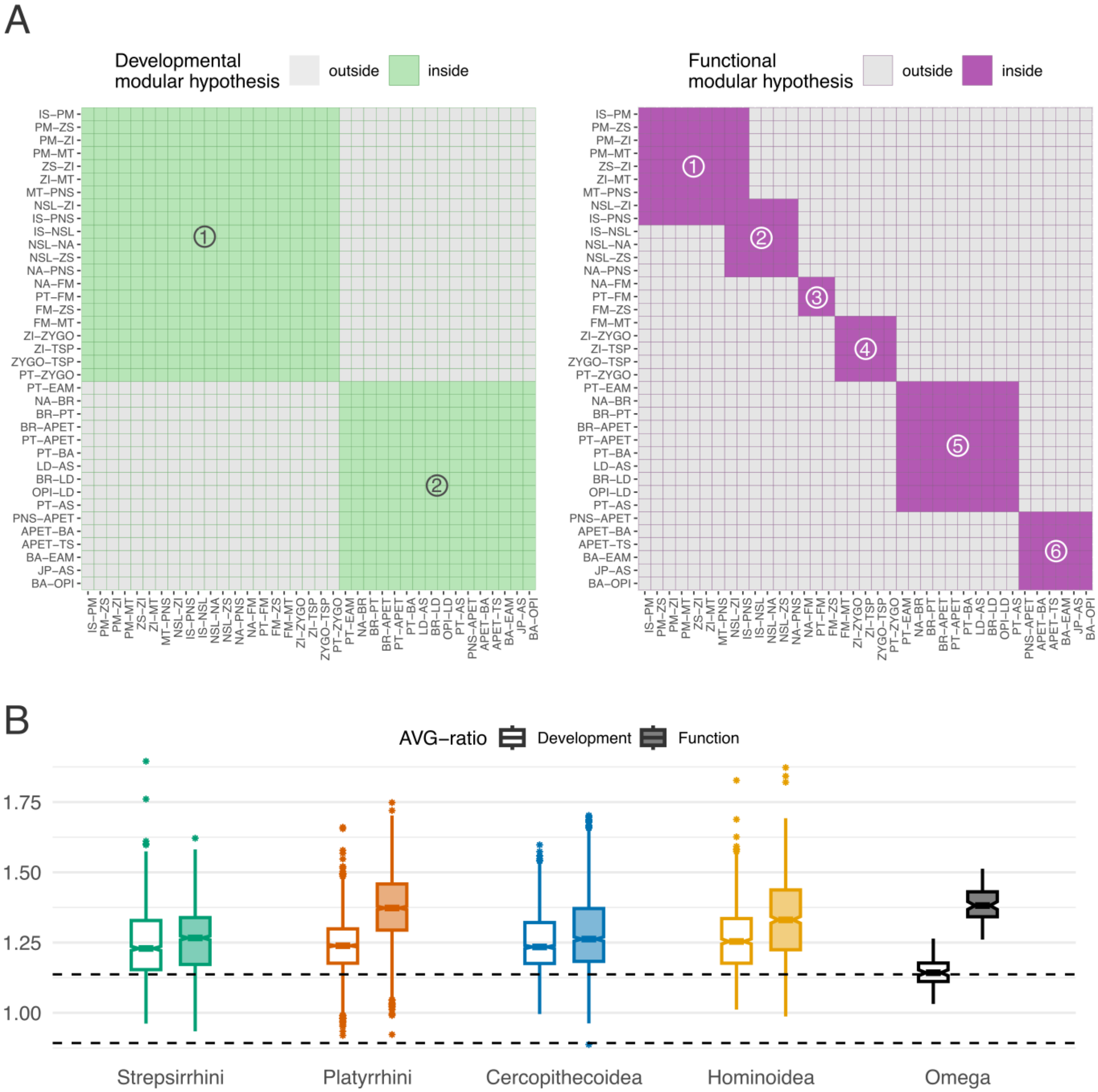


**Fig. S6.** Modularity structure in the primate skull. (**A**) Modular hypothesis tested in this study: Developmental (left), related to the embryonic cell origin of the bones, and Functional (right), related to the functional demands of the enclosing soft tissue throughout life (31–34). These modules can be organized into subunits as: developmental (within-module traits shown in green: 1- rostrofacial, mostly derived from neural crest cells; 2- neurocranium, from the paraxial mesoderm) and functional (within-module traits shown in purple: 1- oral; 2- nasal; 3- orbit; 4- zygomatic; 5- cranial vault; 6- cranial base). For more details about the definition of these modular sets, see (30). In R, these modularity hypothesis matrices were coded as binary matrices, where each row and column correspond to a trait. If the trait in row i is in the same module as the trait in column j, position (i, j) in the modularity hypothesis matrix is set to one; if these traits are not in the same module, position (i, j) is set to zero. (**B**) Boxplots showing the distribution of AVG ratios for the different modular hypothesis tests by clade (colored), and in $\boldsymbol{\Omega}$ (in black), the covariance matrix of all selection gradients. The AVG ratio is computed as the average correlation between traits within the same module and those outside the module for each hypothesis. The area between dashed lines represents the interval of values from the null expectation, obtained by scrambling the rows and columns of the correlation matrices under test.

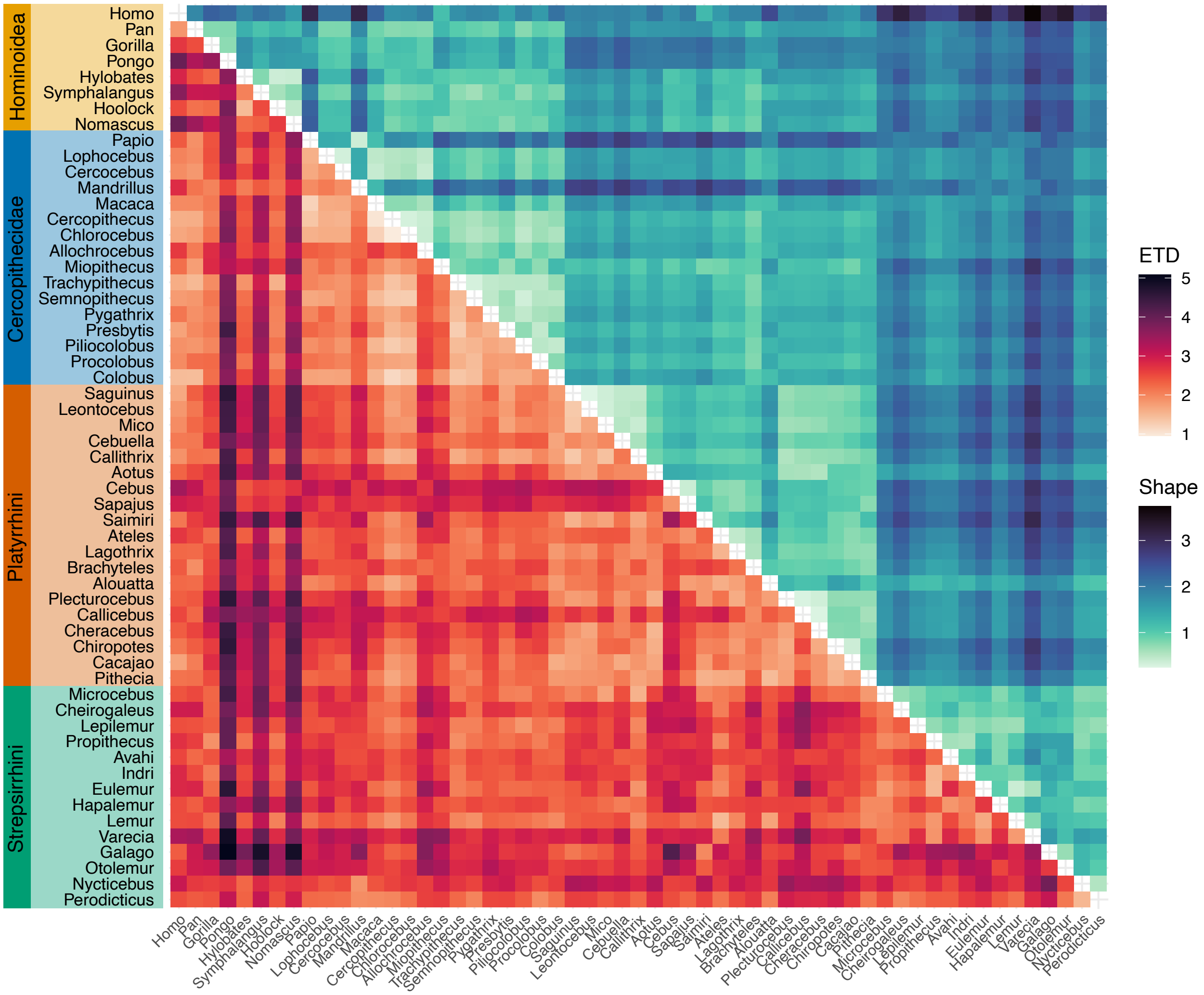


**Fig. S7.** Matrix and morphological divergence are not tightly coupled. Tiles show pairwise distances between genera, with darker tones indicating greater divergence. Distance values for the matrix space were obtained by calculating the Riemannian distances between mean covariance matrices in the Eigen Tensor Decomposition analysis (lower triangle in warm color palette; these values correspond to the y-axis in main text Fig. 3.) Morphological divergence was calculated as the Mahalanobis distances between mean shapes (upper triangle in cold color palette; these values correspond to the x-axis of main text Fig. 3). Morphological divergence shows a broader spread of darker tones than covariance divergence, suggesting that shape has diverged more extensively than covariance structure. This reinforces the idea of a relative stability of integration patterns despite major morphological shifts. Closely related genera (within the same clade) tend to show lighter colors in upper and lower triangles, reflecting lower divergence in both morphology and covariance structure, consistent with phylogenetic signal. More distantly related clades (e.g., Strepsirrhines vs. Hominoids) exhibit consistently darker shades, particularly in morphology, highlighting deep-time divergence. Hominoidea concentrates most of the genera with consistently higher matrix disparity with other primate genera: *Pongo*, *Symphalangus*, and *Nomascus*, reflecting the result obtained from the ETD analysis (Fig. S3).

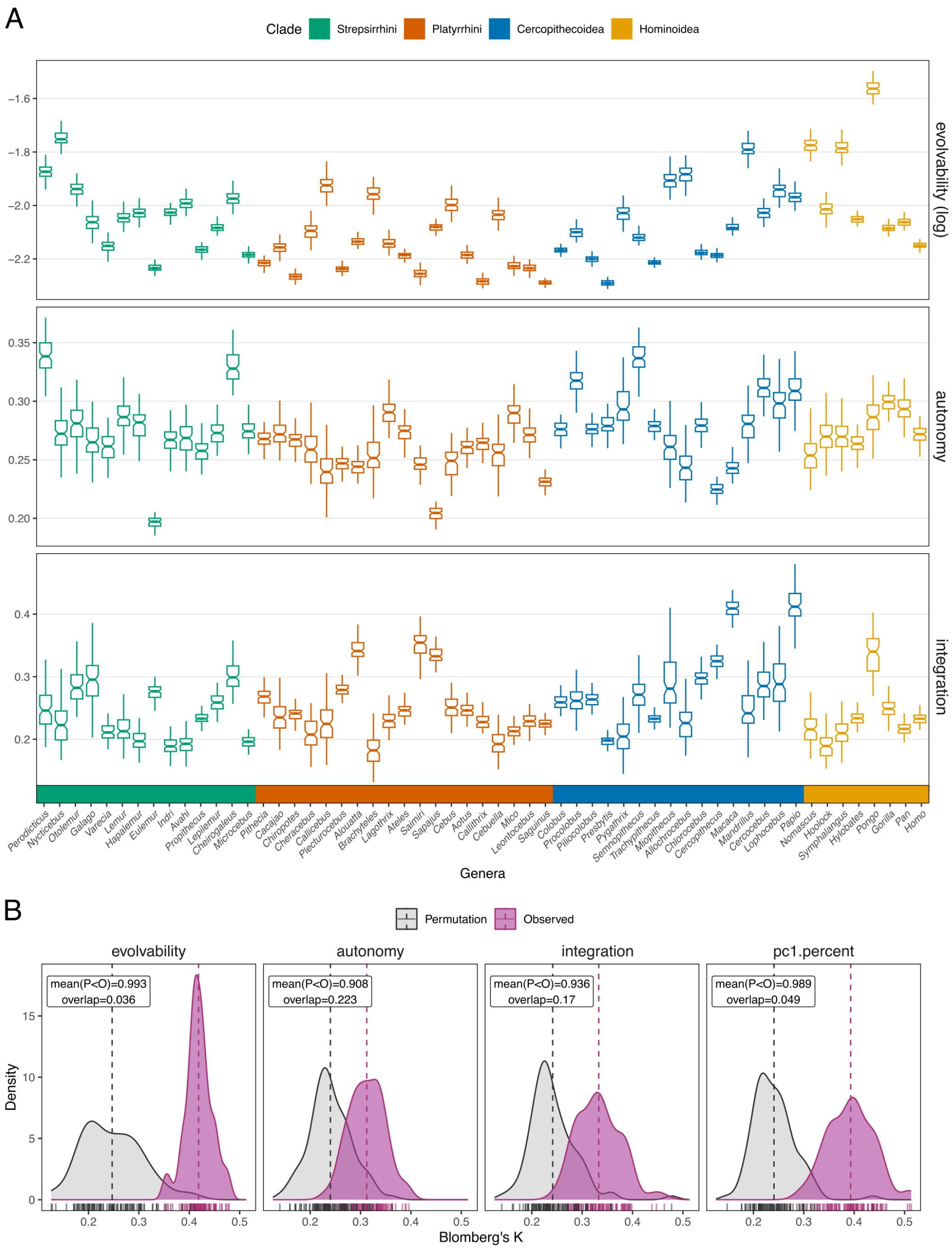


**Fig. S8.** Primates show conserved evolutionary potential and structure of variance. **(A)** Posterior distribution of key evolutionary statistics for each primate genus. Boxplots summarize estimates from 100 posterior phenotypic covariance matrices ($\mathbf{P}$) per genus, showing median values (center line), interquartile ranges (boxes), and whiskers extending to 1.5 times the interquartile ranges. Metrics include overall evolvability, autonomy, and integration, which together describe the magnitude and structure of phenotypic (co)variation relevant to evolutionary potential. Integration (computed as the eigenvalue variance) showed a strong positive correlation with the proportion of variance explained by PC1 (Pearson correlation $= 0.96$, $p < 0.001$). Genera are grouped by major primate clades (Strepsirrhini, Platyrrhini, Cercopithecidea, and Hominoidea; different colors) and sorted phylogenetically. While a few genera diverge from the overall clade average (e.g., *Pongo* for evolvability, *Sapajus* and *Eulemur* for autonomy, and *Papio*, and *Macaca* for integration), there is no clear pattern for divergences. **(B)** Phylogenetic signal for the matrix statistics shown in A and proportion of variance associated with PC1 (Fig. S5. Phylogenetic signal was calculated as Bloomberg's K (which estimates the degree of variation between and within clades). Phylogenetic signal was negligible for autonomy and integration, and moderate for evolvability and PC1 percent. Although observed K values for evolvability and PC1 were less conserved than a pure Brownian Motion expectation (always $< 1$, suggesting greater variance within clades), they were consistently higher than the null distributions obtained by permuting tip labels across all four metrics. This result is strong evidence that closely related genera have more similar matrix statistics than expected by random chance. To account for uncertainty, Blomberg's K was computed using the distribution of matrix statistics calculated from 100 draws from the posterior distributions of P-matrices (Observed values). We then computed a null distribution for the phylogenetic signal by permuting the tip labels, and for each observed value, we calculated the proportion of permutation values lower than or equal to it. Annotation boxes in the top-left corner indicate the mean of the proportion of observations higher than the permuted values, and the proportion of overlapping area between the two distributions (computed as the integral of the minimum between two densities divided by the integral of the maximum of the two densities using the package overlapping in R (36)).

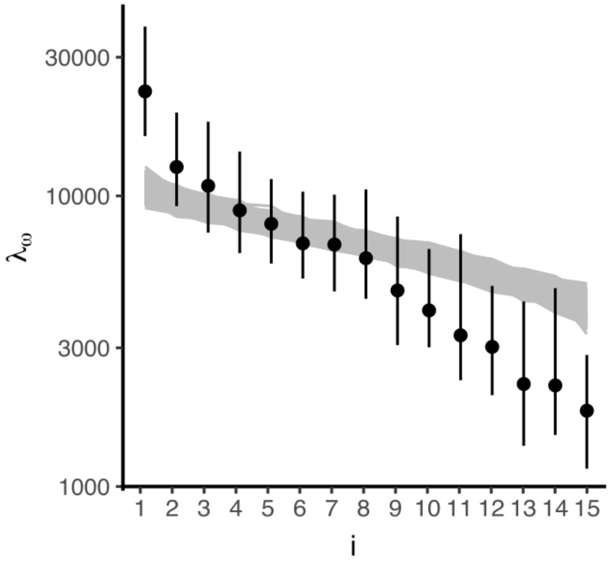


**Fig. S9.** Eccentricity in the distribution of reconstructed selection gradients. Comparison of the empirical eigenspectrum of $\Omega$ with the null expectation of isotropic selection. The black dots represent median values, and vertical bars the $95\%$ credibility intervals. Gray curves correspond to individual simulations under the isotropic null model, in which all directions have the same variance. The observed leading eigenvalues substantially exceed null expectations, demonstrating that reconstructed selection gradients are concentrated along a limited number of preferred directions rather than being uniformly distributed throughout trait space, while the last ones $(12 - 15)$ have less than what would be expected if $\Omega$ were fully isotropic.

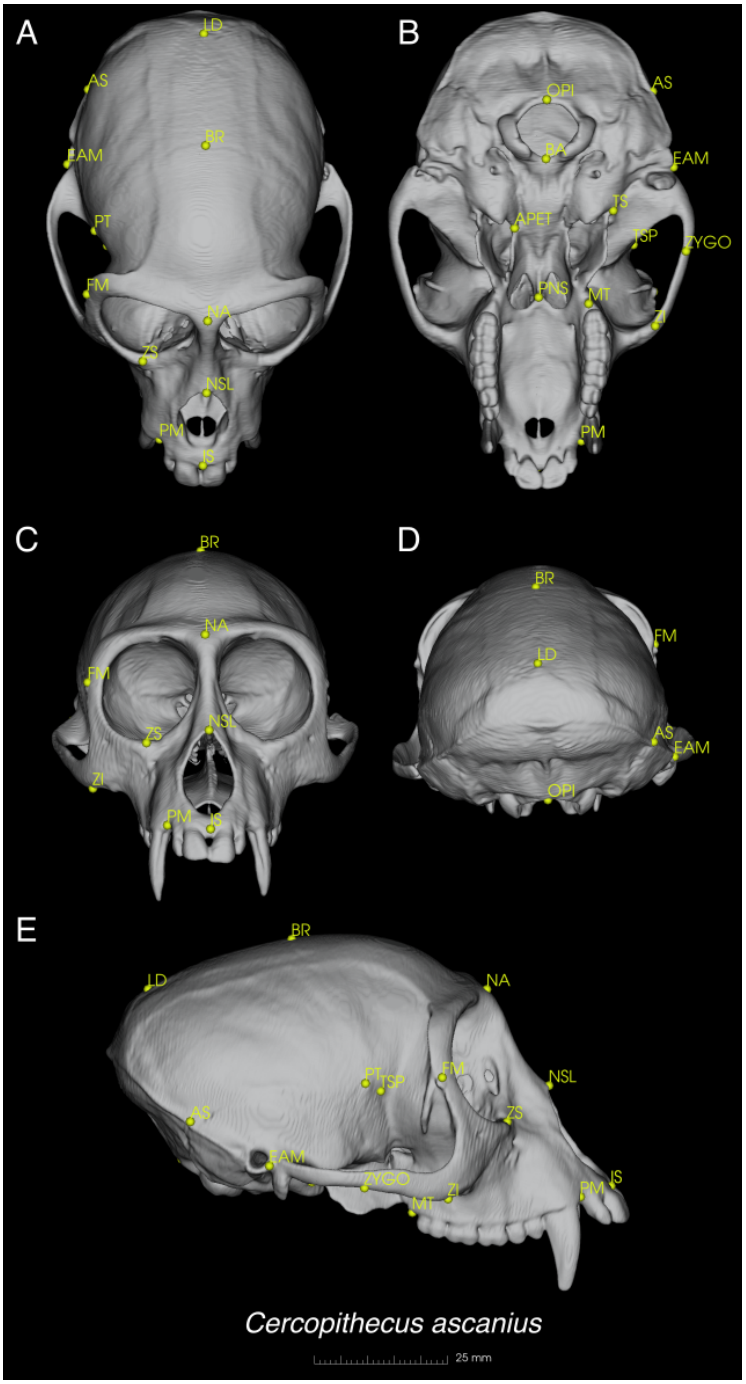


**Fig. S10.** 3D cranial landmarks (in yellow) used in this study exemplified in a specimen of *Cercopithecus ascanius* in A- dorsal, B- ventral, C- anterior, D- posterior, and E- lateral views. A full description of landmarks can be found in Table S1. Image from a CT-scanned specimen available on Morphosource (media ID 000022475,The files was downloaded from www.MorphoSource.org, Duke University).

### Other supporting files

#### SI Dataset S1 (Primaset.RData)

RData file containing a series of objects to replicate the study. Set of 100 matrices per genus, sampled from the posterior distribution of the Bayesian estimation of phenotypic covariance matrices; Median P-matrices for 57 genera; Trait averages for all species, distribution of phylogenetic trees. For a complete description of the dataset, see Primaset_vignette.pdf

#### SI Dataset S2 (Matrix_summary_by_genus.pdf)

Summary of P-matrices, sample size, and trait distributions.
